# In-silico cell sorting revealed granulocyte-specific single-cell-type gene expression from peripheral blood bulk expression data and its application as host response biomarkers to discriminate bacterial and viral infections

**DOI:** 10.64898/2026.04.09.717385

**Authors:** Nelson LS Tang, Tsz-Ki Kwan, Jinghan Huang, Michael LH Tang, Xingyan Wang, Junyi Wu, Christopher KC Lai, Grace CY Lui, Suk-Ling Ma, Kwong-Sak Leung

**Affiliations:** Department of Chemical Pathology, and Li Ka Shing Institute of Health Science, Faculty of Medicine, The Chinese University of Hong Kong, Hong Kong SAR, China; Cytomics Limited, Hong Kong Science Park, Hong Kong SAR, China; Hong Kong Branch of CAS Center for Excellence in Animal Evolution and Genetics and KIZ/CUHK Joint Laboratory of Bioresources and Molecular Research in Common Diseases, Hong Kong SAR, China; Functional Genomics and Biostatistical Computing Laboratory, CUHK Shenzhen Research Institute, Shenzhen, China; Beijing Anzhen Hospital, Capital Medical University, Beijing, China; Department of Microbiology, Faculty of Medicine, The Chinese University of Hong Kong, Hong Kong SAR, China; Department of Medicine & Therapeutics, Faculty of Medicine, The Chinese University of Hong Kong, Hong Kong SAR, China; School of Arts and Humanities, Tung Wah College, Hong Kong SAR, China; Department of Computer Science and Engineering, Faculty of Engineering, The Chinese University of Hong Kong, Hong Kong SAR, China

## Abstract

Peripheral Blood transcriptome analysis evaluated the bulk transcript abundance (TA) covering all leukocyte cell populations. However, there are 2 main problems in using bulk expression as biomarkers: (1) A long list of differential expression genes (DEGs) was found, and (2) DEGs cannot be attributed to a host response of any specific cell-type. TA assays after conventional cell sorting, as the gold-standard method, is too tedious for routine use. Recently, we showed that by using a ratio-based biomarker, RBB (ratio of two stringently selected genes), it is feasible to interrogate the gene expression of a single cell-type (monocyte and B lymphocyte) in peripheral whole blood (WB) directly. Here, we apply this in-silico cell sorting algorithm (DIRECT LS-TA, Direct Leukocyte Single cell-type Transcript Abundance) to granulocytes in WB samples to reveal RBBs specific to granulocytes. This DIRECT LS-TA approach without the need for cell-sorting was applied to public datasets to differentiate the 2 types of infection (bacterial vs viral infection). The following RBBs measured in WB correlate with the expression of target (numerator) genes in purified granulocytes, thus cell-sorting can be avoided by using these RBBs: *ARG1/SRGN, ANXA3/SRGN, RSAD2/SRGN*. Together with monocyte DIRECT LS-TA biomarkers*, IFI27/PSAP,* direct quantification of 4 genes provided optimal differentiation of viral from bacterial infection. Meta-analysis and unsupervised machine learning classification confirmed the superior performance of DIRECT LS-TA biomarkers. These RBBs found by prior In-silico cell-sorting identified pairs of genes that are used to formulate as ratio-based biomarkers (RBBs) to represent gene expression of granulocytes inside whole blood cell-mixture samples which was useful to triage febrile patients into two major categories of febrile diseases between viral and bacterial infection with high degree of sensitivity and specificity.

## Introduction

Febrile illness is a common presentation, and rapid triage of patients can significantly improve clinical management. Currently, laboratory methods primarily focus on detecting or culturing pathogens to establish a diagnosis. However, a definitive pathogen was not detected in up to 75% of febrile children even using the latest molecular pathogen tests [1]. So far, little attention has been given to analysing the host response, except for the use of various serum proteins (such as C-reactive protein and procalcitonin) to reflect the extent of ongoing inflammation in a non-specific manner. However, in the past five years, there has been a new initiative of using blood cell transcriptome to differentiate viral from bacterial infections. Last year, a large-scale validation study showed that a host response gene expression blood test can discriminate bacterial from viral infection with high sensitivity and specificity of over 80% [2]. Similar approaches of using host transcriptome to differentiate the cause of infection are reviewed recently [3,4]. However, most of them involve the quantification of large number of genes and is only available from specialized laboratories, which limits its wider application [2,5–7].

Buonsenso et al recently reviewed studies on the host blood transcriptome in febrile children [4]. These studies commonly employed transcriptome profiling methods such as microarrays and bulk RNA-seq to identify differentially expressed genes (DEG) between groups of patients with viral infections versus bacterial infections or other illnesses [8,9]. However, they all used standard normalisation and DEG analysis approaches, which typically generated a long list of DEGs. Consequently, tens to hundreds of DEGs were used as classifier signatures [5,10,11]. There are only a few exceptional example gene panels requiring fewer DEGs as biomarkers [12–18]. Most of these exceptions used interferon-stimulated genes (ISGs, such as *IFI44L*) as biomarkers to diagnose viral infections, which induce an interferon response. However, these small gene panels may not be able to differentiate other febrile conditions from signature gene panels comprising more genes.

There are two major shortcomings in using the conventional DEG approach. Firstly, it is unclear which leukocyte cell-type(s) in the blood are responsible for the gene activation response. As blood is a mixture of various leukocyte cell-types, it would be more informative to know which specific cell type is responsible for the activation. Typically, this would require single-cell RNAseq (scRNA-seq) analysis [19]. However, the cost and sample requirements of scRNA-seq make it not feasible for large-scale clinical applications. A simple method for estimating the appropriate level of gene expression in a specified cell population in blood would be very useful.

To address these issues, we introduced a rapid method to directly measure gene expression in single cell types from blood samples, called the <u>D</u>irect <u>L</u>eukocyte <u>S</u>ingle cell-type <u>T</u>ranscript <u>A</u>bundance (DIRECT LS-TA) assay, which recently targets monocytes [20,21]. The assay utilises a small panel of only a few genes (as few as three precisely selected monocyte-informative genes) and can distinguish between viral and bacterial infections. These monocyte-informative genes are chosen through an algorithm known as in silico cell-sorting, which identifies genes in peripheral blood that are predominantly expressed by monocytes. Single-cell-type-informative genes can be identified from transcriptome data using ICEBERG plots. The ratio of two monocyte-specific genes serves as a ratio-based biomarker (RBB), called DIRECT LS-TA. For example, the *IFI27/PSAP* ratio serves as a monocyte-specific biomarker of interferon activation and provides an excellent indicator of the host response to viral infection.

### In-silico cell-sorting and a simple ratio-based biomarker (RBB) approach can estimate the single cell-type specific gene expression from bulk (cell mixture) samples - DIRECT LS-TA assay

Most blood transcriptome data available in the public domain are bulk gene expression data that represent the summation of transcripts expressed by all cells in a cell-mixture sample composed of various populations of specific cell types (such as T-cells, monocytes, granulocytes, etc.). The major drawback of analyzing bulk expression data is the inability to determine if the change in expression is due to a change in cell number or if it represents gene activation in a particular cell type. Only the latter is of biological interest and would serve as a robust biomarker. Therefore, various approaches have been developed to estimate gene expression in specific cell populations or cell counts from bulk gene expression data. The first approach was based on mathematical deconvolution [22], which has become the foundation of most other methods available today. A more widely used package is CIBERSORT [23], which has shown favourable performance when benchmarked against other algorithms [24]. However, transcriptome-scale data are required for deconvolution to function. Therefore, it cannot be applied in a clinical setting due to both financial constraints and turnaround time.

We pioneered a new method, based on given cell count proportions of various leukocyte single-cell-types in peripheral blood and differential expression levels of genes in each of these cell-types, directly estimated expression levels of some selected genes for a single cell-type (e.g. B cell [25], or monocytes [20]) in the blood samples without cell sorting. The purpose of in-silico cell-sorting is to shortlist genes that are predominantly expressed by only one single cell-type in a cell-mixture sample (e.g. WB). The basic idea is, for a specified leukocyte cell-type, to define a list of genes that are highly expressed by this leukocyte population so that the majority of mRNA transcripts in the blood sample originated from this specified single cell-type (for example, granulocytes as the cell-type of interest here). And these genes are called cell-type-informative genes for granulocytes, for example. As granulocytes contributed the majority of transcripts of these genes in WB, when TA levels of these genes are assayed in WB, it reflects to a very large extent the expression in granulocytes. In other words, when TA of granulocyte-informative genes are measured in peripheral blood, the results will be highly correlated with those expressed in the purified granulocytes collected after cell sorting, which is taken as the gold standard. In our previous manuscript, we showcased the application to monocytes, and a list of about 100 monocyte-informative genes has been reported [20].

Figure 1 explains how this in-silico cell-sorting algorithm works. Then, the next step is to obtain a robust and reproducible biomarker representing TA of a target gene expressed in individual granulocytes in the samples. We showed that single-cell TA (or TA per granulocytes) for some target genes was related to a ratio (RBB) using two granulocyte-informative genes: one target gene as the numerator and another as the denominator in the RBB. The target gene is responsive to stimulations of interest. The denominator is another granulocyte-informative gene, but it also has the least biological variation (e.g., between-individual variance). This in-silico cell sorting model predicts that the RBB (simply a ratio of expression levels of 2 genes measured in WB, correlates with expression of the target gene in the specified single cell-type purified by cell sorting. We showed how it worked for monocytes to detect host response to bacterial infection [20]. This DIRECT LS-TA approach eliminates the tedious or expensive procedures of manual cell sorting or scRNA-seq. Such in-silico cell sorting algorithm can be done using selected public gene expression datasets (e.g. in GEO database) to obtain a list of cell-type informative genes.

**Figure 1.**
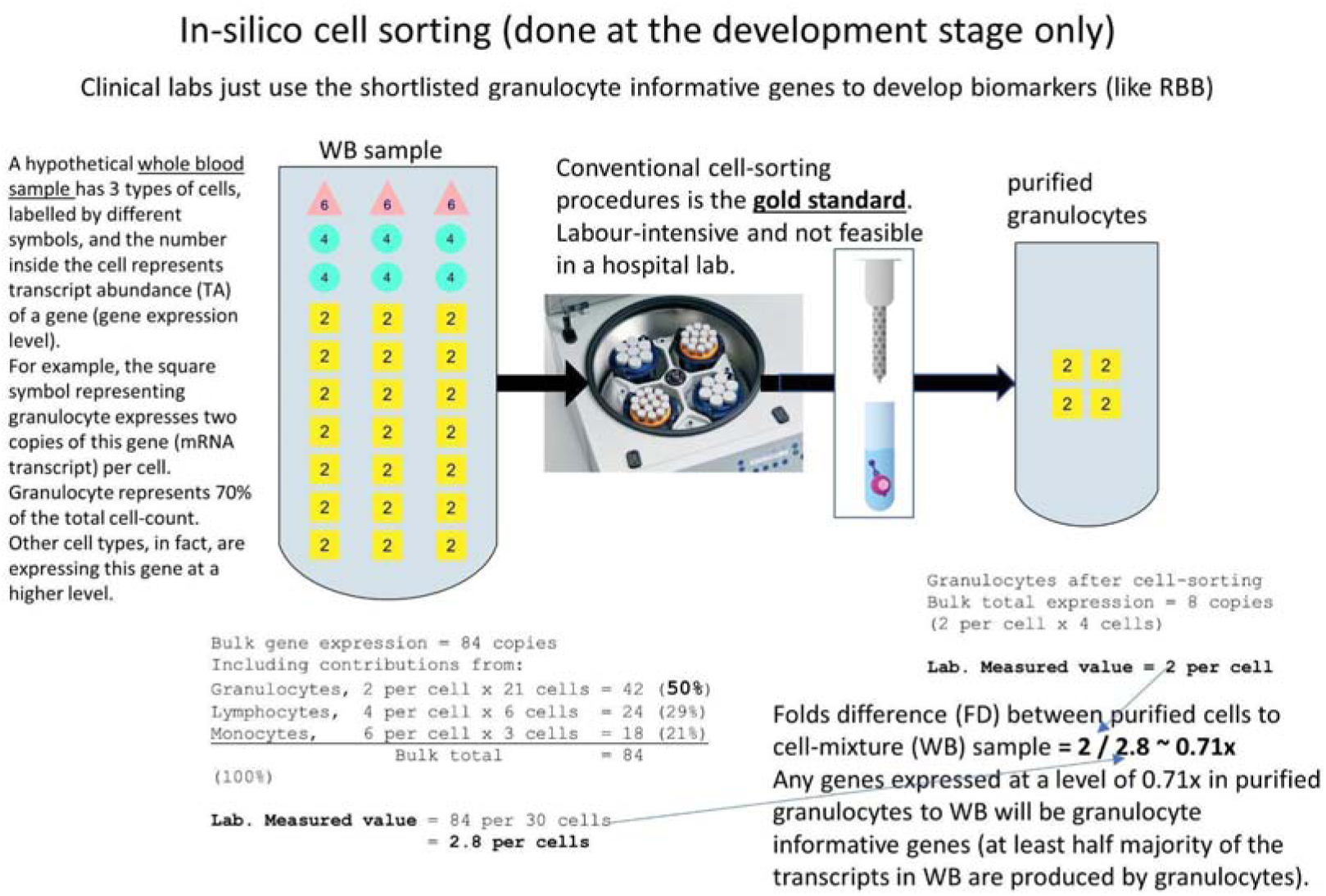
A schematic diagram to illustrate the concept underlying in-silico cell-sorting to define single-cell-type (granulocyte) informative genes in WB samples. WB collected in the sample tube contains various cell populations, for simplicity 3 cell-types are shown here by different shape symbols. Circles represent granulocytes accounting for 70% of all cells. Squares represent lymphocytes and triangles represent monocytes. The numbers inside the symbol represent a hypothetical per-cell expression level for one gene. Each granulocyte produces 2 copies per cell, lymphocyte: 4 and monocyte: 6. Even granulocytes’ per cell production of this gene transcript is lower than both lymphocytes and monocytes. Granulocyte contribute the majority of the transcripts of the gene in the bulk WB sample due to its high proportional cell-count. As shown, granulocytes produce 42 of 84 transcripts. It reaches the definition of a single-cell-type informative gene. Single-cell-type informative gene can be revealed by comparing the extent of enrichment (fold difference, FD) of gene expression levels between a purified cell sample and the WB cell-mixture sample. As shown, the FD for this gene is as low as 2/2.8 = 0.71-fold when granulocytes account for 70% of WB cell counts. In our work here, a much higher FD of 2 is used to shortlist granulocyte-informative genes shown in Table 1. Therefore, granulocytes account for the great majority (up to 90%) of mRNA transcripts for the genes listed in Table 1 in WB.

Here, we extend the in-silico cell-sorting algorithm to granulocytes in WB (Figure 1). As granulocytes account for 70% or more of leukocytes in WB, our DIRECT LS-TA approach leverages this high cellular proportion and, for the first time, unambiguously identifies hundreds of granulocyte-informative genes. Therefore, the measurement of expression of these genes directly in WB can be used to reflect single-cell expression levels of granulocytes inside the WB sample.

## Materials and Methods

In this study, we aimed to develop the granulocyte DIRECT LS-TA method to directly obtain TA from granulocytes in peripheral blood samples. The objective is to develop and validate granulocyte-specific DIRECT LS-TA ratio-based biomarkers (RBB) that can be readily implemented in a clinical setting to detect the host response to infectious and inflammatory diseases.

### Public Gene Expression Datasets

To identify granulocyte-informative genes suitable for the DIRECT LS-TA assay, in-silico cell-sorting was applied to the following gene expression datasets. These datasets were available from the Gene Expression Omnibus (GEO), maintained by the US National Institutes of Health. Details were available under their accession numbers.

For the selection of granulocyte-informative genes by the in-silico cell sorting method, datasets used must contain gene expression data of both manually purified granulocytes (or neutrophils) and corresponding whole blood (WB) samples from the same individuals. These paired datasets are also essential for (1) creating the ICEBERG plot to select granulocyte-informative genes and (2) confirming the correlation between the granulocyte-specific DIRECT LS-TA ratio-based biomarker (RBB) of target genes in Table 1 and the gold-standard gene expression results obtained from purified granulocyte samples.

**Table 1.**
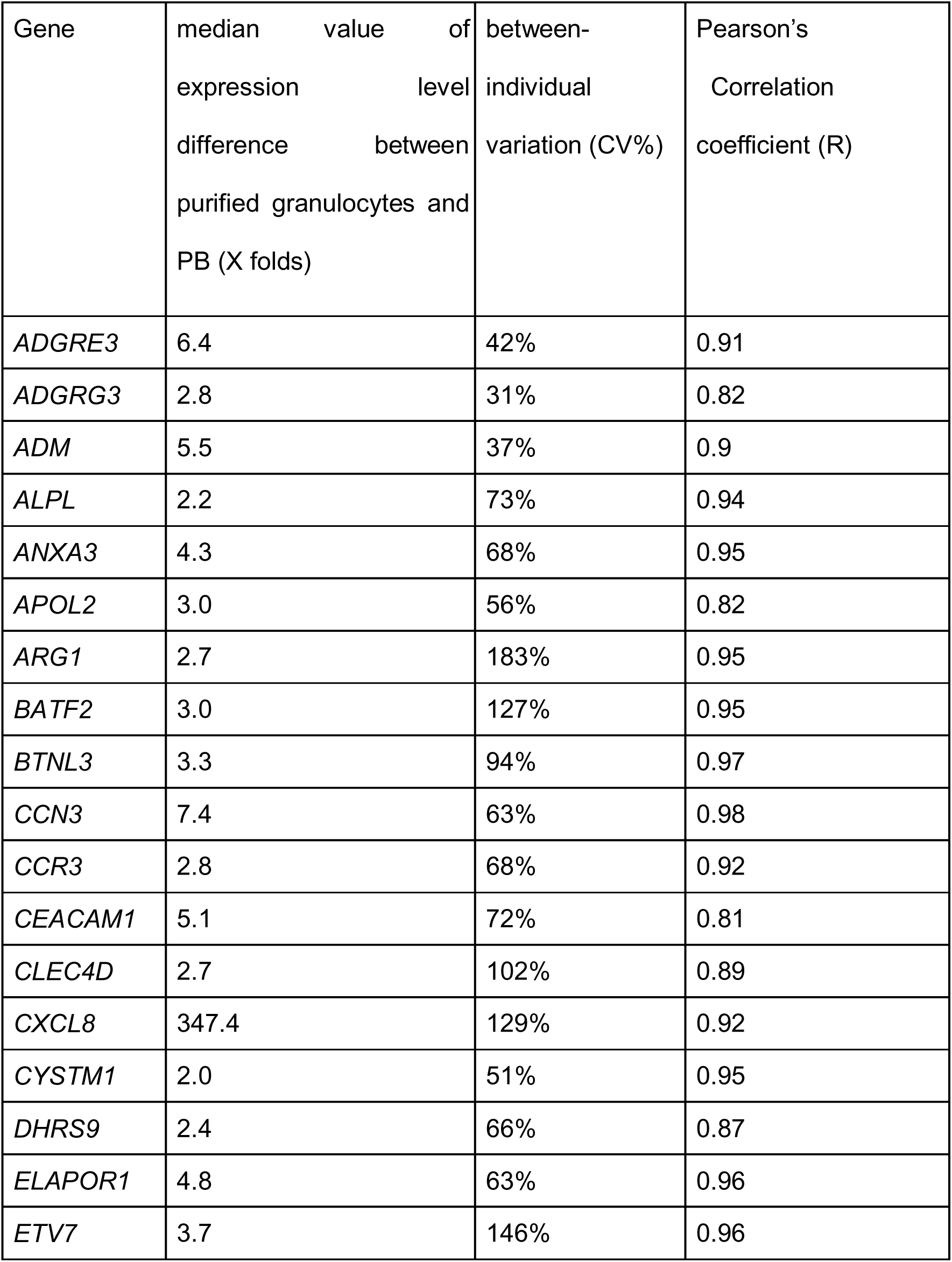
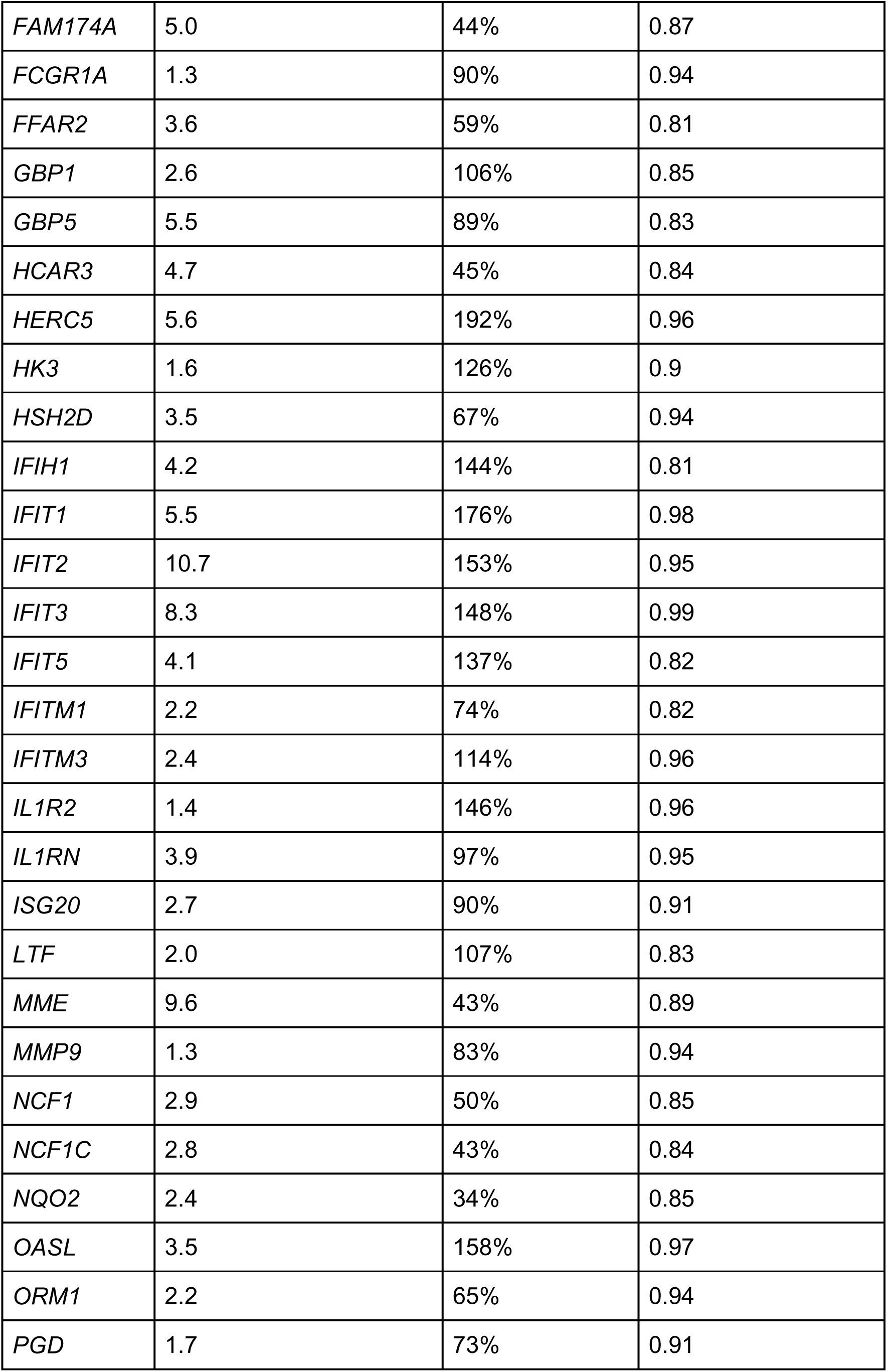
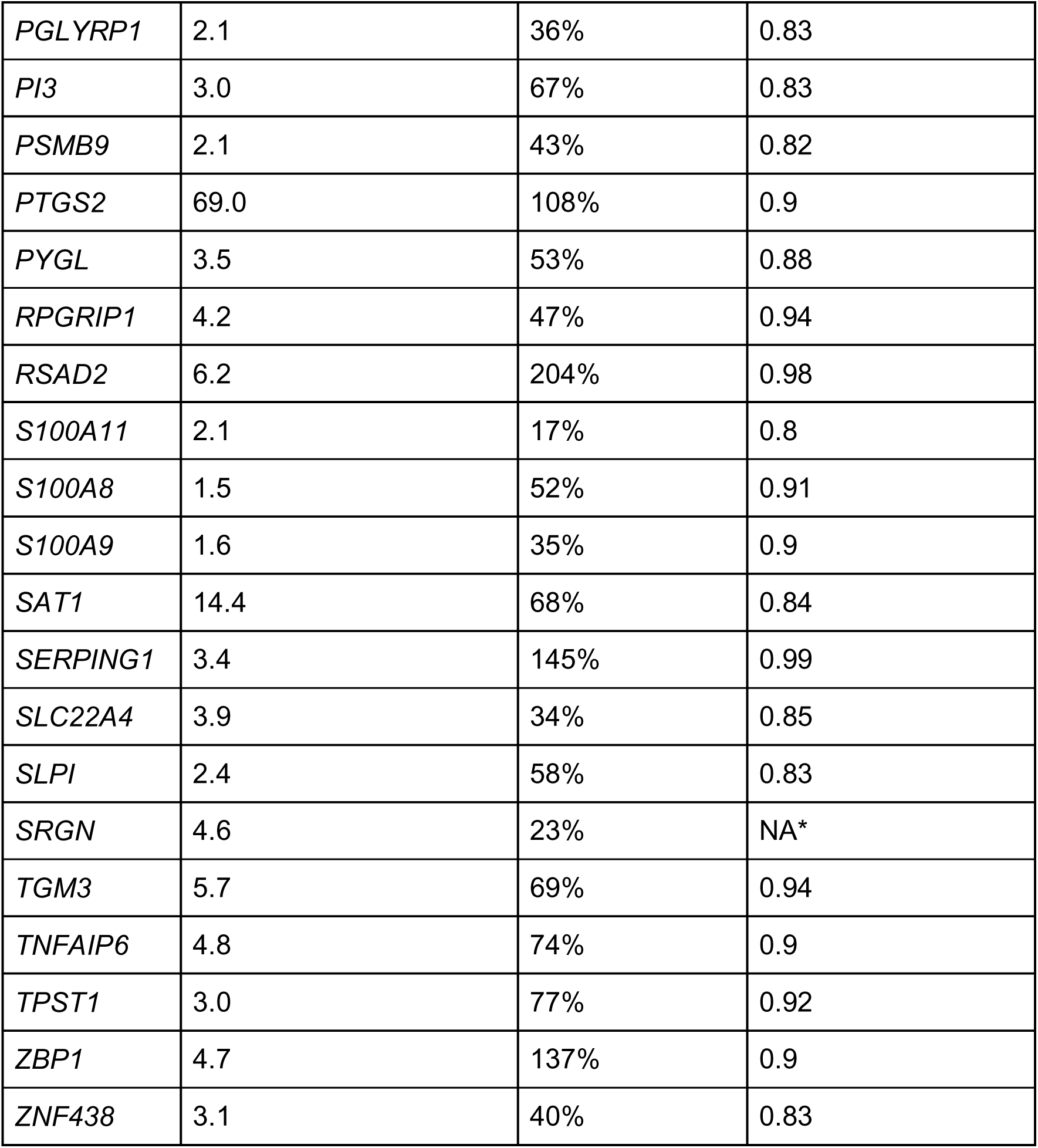
List of granulocyte-informative genes. * *SRGN* is used as denominator gene in the RBB, so it do not have correlation results to compare with gold standard method.

| Gene | median value of expression level difference between purified granulocytes and PB (X folds) | between-individual variation (CV%) | Pearson's Correlation coefficient (R) |
| --- | --- | --- | --- |
| *ADGRE3* | 6.4 | 42% | 0.91 |
| *ADGRG3* | 2.8 | 31% | 0.82 |
| *ADM* | 5.5 | 37% | 0.9 |
| *ALPL* | 2.2 | 73% | 0.94 |
| *ANXA3* | 4.3 | 68% | 0.95 |
| *APOL2* | 3.0 | 56% | 0.82 |
| *ARG1* | 2.7 | 183% | 0.95 |
| *BATF2* | 3.0 | 127% | 0.95 |
| *BTNL3* | 3.3 | 94% | 0.97 |
| *CCN3* | 7.4 | 63% | 0.98 |
| *CCR3* | 2.8 | 68% | 0.92 |
| *CEACAM1* | 5.1 | 72% | 0.81 |
| *CLEC4D* | 2.7 | 102% | 0.89 |
| *CXCL8* | 347.4 | 129% | 0.92 |
| *CYSTM1* | 2.0 | 51% | 0.95 |
| *DHRS9* | 2.4 | 66% | 0.87 |
| *ELAPOR1* | 4.8 | 63% | 0.96 |
| *ETV7* | 3.7 | 146% | 0.96 |
| *FAM174A* | 5.0 | 44% | 0.87 |
| *FCGR1A* | 1.3 | 90% | 0.94 |
| *FFAR2* | 3.6 | 59% | 0.81 |
| *GBP1* | 2.6 | 106% | 0.85 |
| *GBP5* | 5.5 | 89% | 0.83 |
| *HCAR3* | 4.7 | 45% | 0.84 |
| *HERC5* | 5.6 | 192% | 0.96 |
| *HK3* | 1.6 | 126% | 0.9 |
| *HSH2D* | 3.5 | 67% | 0.94 |
| *IFIH1* | 4.2 | 144% | 0.81 |
| *IFIT1* | 5.5 | 176% | 0.98 |
| *IFIT2* | 10.7 | 153% | 0.95 |
| *IFIT3* | 8.3 | 148% | 0.99 |
| *IFIT5* | 4.1 | 137% | 0.82 |
| *IFITM1* | 2.2 | 74% | 0.82 |
| *IFITM3* | 2.4 | 114% | 0.96 |
| *IL1R2* | 1.4 | 146% | 0.96 |
| *IL1RN* | 3.9 | 97% | 0.95 |
| *ISG20* | 2.7 | 90% | 0.91 |
| *LTF* | 2.0 | 107% | 0.83 |
| *MME* | 9.6 | 43% | 0.89 |
| *MMP9* | 1.3 | 83% | 0.94 |
| *NCF1* | 2.9 | 50% | 0.85 |
| *NCF1C* | 2.8 | 43% | 0.84 |
| *NQO2* | 2.4 | 34% | 0.85 |
| *OASL* | 3.5 | 158% | 0.97 |
| *ORM1* | 2.2 | 65% | 0.94 |
| *PGD* | 1.7 | 73% | 0.91 |
| *PGLYRP1* | 2.1 | 36% | 0.83 |
| *PI3* | 3.0 | 67% | 0.83 |
| *PSMB9* | 2.1 | 43% | 0.82 |
| *PTGS2* | 69.0 | 108% | 0.9 |
| *PYGL* | 3.5 | 53% | 0.88 |
| *RPGRIP1* | 4.2 | 47% | 0.94 |
| *RSAD2* | 6.2 | 204% | 0.98 |
| *S100A11* | 2.1 | 17% | 0.8 |
| *S100A8* | 1.5 | 52% | 0.91 |
| *S100A9* | 1.6 | 35% | 0.9 |
| *SAT1* | 14.4 | 68% | 0.84 |
| *SERPING1* | 3.4 | 145% | 0.99 |
| *SLC22A4* | 3.9 | 34% | 0.85 |
| *SLPI* | 2.4 | 58% | 0.83 |
| *SRGN* | 4.6 | 23% | NA* |
| *TGM3* | 5.7 | 69% | 0.94 |
| *TNFAIP6* | 4.8 | 74% | 0.9 |
| *TPST1* | 3.0 | 77% | 0.92 |
| *ZBP1* | 4.7 | 137% | 0.9 |
| *ZNF438* | 3.1 | 40% | 0.83 |

For validation of clinical utility of various DIRECT LS-TA RBBs, other datasets from clinical cohorts were used. To differentiate bacterial from viral infections, for example, a discovery dataset was used to identify potential granulocyte DIRECT LS-TA RBBs activated in granulocytes after bacterial infection, and these RBBs were then evaluated across multiple replication datasets. Different gene expression platforms were included here to show that the new granulocyte DIRECT LS-TA was a genuine granulocyte phenomenon and not analytical platform-dependent.

### In-silico cell-sorting: Step 1 Identification of Granulocyte-Informative Genes

Single-cell-type informative genes is defined as any gene with the majority (at least 50%) of its mRNA transcripts in a cell-mixture that is produced by a single leukocyte cell-type. The selection of candidate granulocyte-type informative genes suitable for granulocyte-type informative expression analysis in cell-mixture samples is based on differences in expression levels (fold difference, FD) between the manually purified cell-fraction sample of the specified cell type and the cell-mixture sample. Genes meeting this FD (also called X50 criteria, meaning FD at 50% contribution from a single cell type) were shortlisted as candidate granulocyte single-cell-type-informative genes. This is the mathematical basis of DIRECT LS-TA assays and the ICEBERG plot [20]. An ICEBERG plot shows the FD values of all genes on the y-axis. A pre-defined FD cut-off value, for example, two folds, can be used to select single-cell type informative genes with FD greater than 2, resembling the part of the ICEBERG that can be seen above the sea level, while the greater part of the ICEBERG (other non-informative genes) is submerged in the sea and cannot be seen.

The fold difference (X50) required for granulocyte-informative genes was calculated using the formula X = 1/(2P). For WB samples with 70% granulocyte cell count as shown in Figure 1 (granulocytes with P = 0.7), the required FD X50 is only 0.71-fold. We selected a much more stringent cut-off value of FD=1, so that shortlisted genes are largely contributed by granulocytes alone in WB. So that we can reliably determine their granulocyte-specific expression directionally in WB by using Direct LS-TA RBB.

### In-silico cell-sorting: Step 2 Selection of Reference and Target Genes

The ICEBERG plot is a scatter plot showing the differential TA of ∼20,000 genes between purified granulocytes and WB (FD) as the Y axis. FD can be plotted against either gene expression level (TA) or biological variation (e.g., between-individual variance). Only those genes with differential expression (FD, Y-axis values) greater than 1 are potential granulocyte-informative genes.

Two types of granulocyte-informative genes are needed to construct the granulocyte DIRECT LS-TA RBB:

1. Granulocyte-informative **reference genes** are selected from the single-cell-type-informative genes identified for granulocytes. They had low-level biological variation. They serve as the denominator in the DIRECT LS-TA RBB and were selected for having the lowest biological variation, as measured by the geometric coefficient of variation (CV)
2. Granulocyte-informative **target genes** (also referred to as single-cell-type target genes): These genes may be involved in a pathway of interest and may be differentially expressed in healthy subjects and patients. The granulocyte-informative target gene serves as the numerator of the DIRECT LS-TA RBB.

### DIRECT LS-TA RBB: the biomarker is simply a ratio of 2 selected genes

The new RBB is a ratio of TA of 2 genes measured directly in the cell-mixture sample (e.g. WB):

Granulocyte DIRECT LS-TA = TA of granulocyte-informative target gene (WB)/ TA of granulocyte-informative reference gene (WB)

In the gene expression dataset, the gene expression values are log-transformed. Therefore, this RBB can also take its log form as follows:

Log granulocyte DIRECT LS-TA = Log TA of granulocyte-informative target gene (WB) - Log TA of granulocyte-informative reference gene (WB)

Conceptually, this is similar to the results of delta CT (ΔCT) in quantitative PCR (qPCR) relative quantification experiments. The difference of threshold cycles (CT) of the target gene and the normalisation reference gene (typically one or more housekeeping genes) is called delta CT (ΔCT) in qPCR relative quantification. Direct LS-TA RBB can also be expressed as delta-CT when used in qPCR assays but the typical housekeeping gene is replaced by a granulocyte informative reference gene mentioned in the previous sections.

### Statistical Analysis

Group-wise differences in granulocyte DIRECT LS-TA results were compared between the control group and the patient group using non-parametric statistical tests (e.g. Wilcoxon-Mann-Whitney test). A multiple testing correction by the Bonferroni method was used where appropriate, and the type I error is set to the corrected significance level. The diagnostic performance of granulocyte DIRECT LS-TA biomarkers was evaluated using receiver-operating characteristic (ROC) analysis, and the area-under-curve (AUC) was calculated to quantify discriminative performance.

## Result

### In silico cell sorting: identify granulocyte-informative genes by ICEBERG plot

GSE60424 was used to identify granulocyte-informative genes because it included data from both WB and purified granulocytes. The y-axis of the ICEBERG plot shows the differential expression FD (folds as unit) between purified granulocytes and the corresponding WB sample. The higher this FD value, the higher the contribution of granulocytes to the sum of mRNA transcripts of the gene in WB, or in other words, the transcript of this gene is more specifically originated from granulocytes. Based on the ICEBERG plot (Figure 2A and 2B), genes above this cut-off value of FD=1x are defined as granulocyte-informative genes.

**Figure 2.**
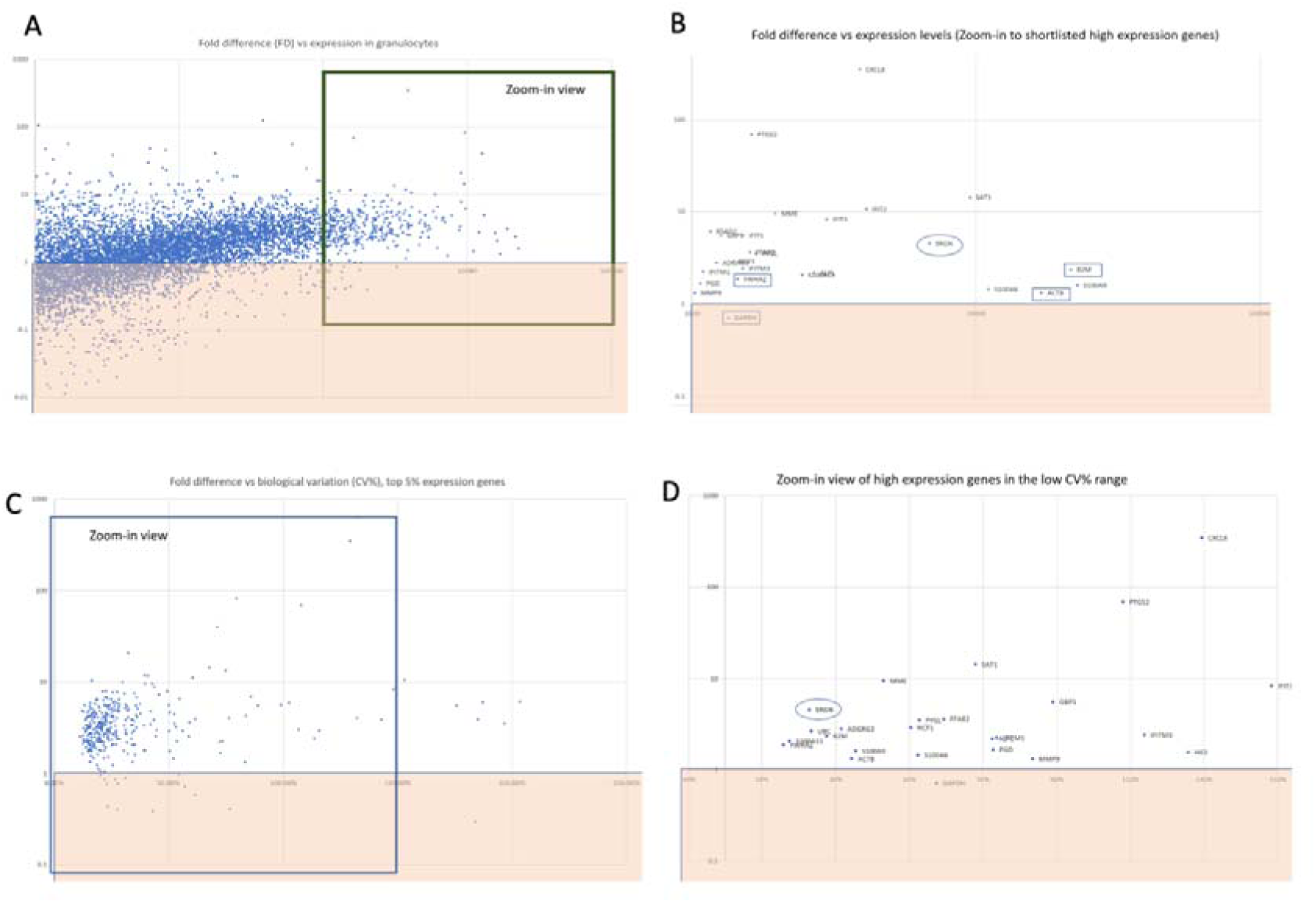
The ICEBERG plot. (A) ICEBERG plot with Fold Difference (x-fold) indicating mRNA transcript enrichment in granulocytes over WB, as well as the expression levels of genes. (B) ICEBERG plot of shortlisted genes. (C) ICEBERG plot with fold Difference (x folds) as an indication of mRNA transcript enrichment in granulocytes over WB vs biological variation of expression (CV%). In order to search for granulocyte-informative reference genes to be used as the denominator in the DIRECT LS-TA RBB, the region of low CV% is zoomed in to show in Figure 2D. (D) ICEBERG plot of shortlisted genes with low CV%. SRGN (marked by an oval) is selected as a granulocyte-informative reference gene because it has a high FD value and a low CV%. Other typical housekeeping genes are also shown for comparison (e.g. ACTB, B2M, GAPDH), which also have similar low biological variation CV%. However, they are below sea level, as they have FD values much lower than the cut-off value (c.f. SRGN or SAT1).

In fact, almost half of the genome (up to 10,000 genes) is expressed above this cut-off value, Figure 2A, confirming that TA of most genes in WB predominantly reflects the expression of granulocytes. We then confined to those high-expression genes that had TA in the top 15% of all genes (Figure 2B).

Two genes, *SRGN* and *SAT1,* were selected as granulocyte-informative reference gene, as they are both abundantly produced by granulocytes in WB and they had the least biological variation (measured as between-individual variation) (Figure 2C and 2D). They will be used as denominators to derive ratio-based biomarkers (RBB) in subsequent biomarker evaluation. Although many genes had fold-value X of greater than 1, we confined to those with excellent correlation (R>0.7) between DIRECT LS-TA and gene expression in manually purified granulocytes for subsequent analysis. The shortlisted granulocyte-informative genes are shown in Table 1, along with their correlations with gene expression in purified granulocytes.

### In-silico cell-sorting validation by excellent correlation between DIRECT LS-TA RBB in WB and gene expression in manually purified granulocyte

A validation step was taken to confirm that Direct LS-TA RBB can represent gene expression in manually purified granulocytes. All granulocyte-informative genes listed in Table 1 showed a high correlation between DIRECT LS-TA RBB in WB (the ratio of the target gene to *SRGN* measured in WB) to target gene expression (normalised to a typical housekeeping gene, *UBC*) in manually purified granulocytes. All granulocyte-informative genes shown in the table had correlation coefficients greater than 0.8 (all statistically significant) (Figure 3).

**Figure 3.**
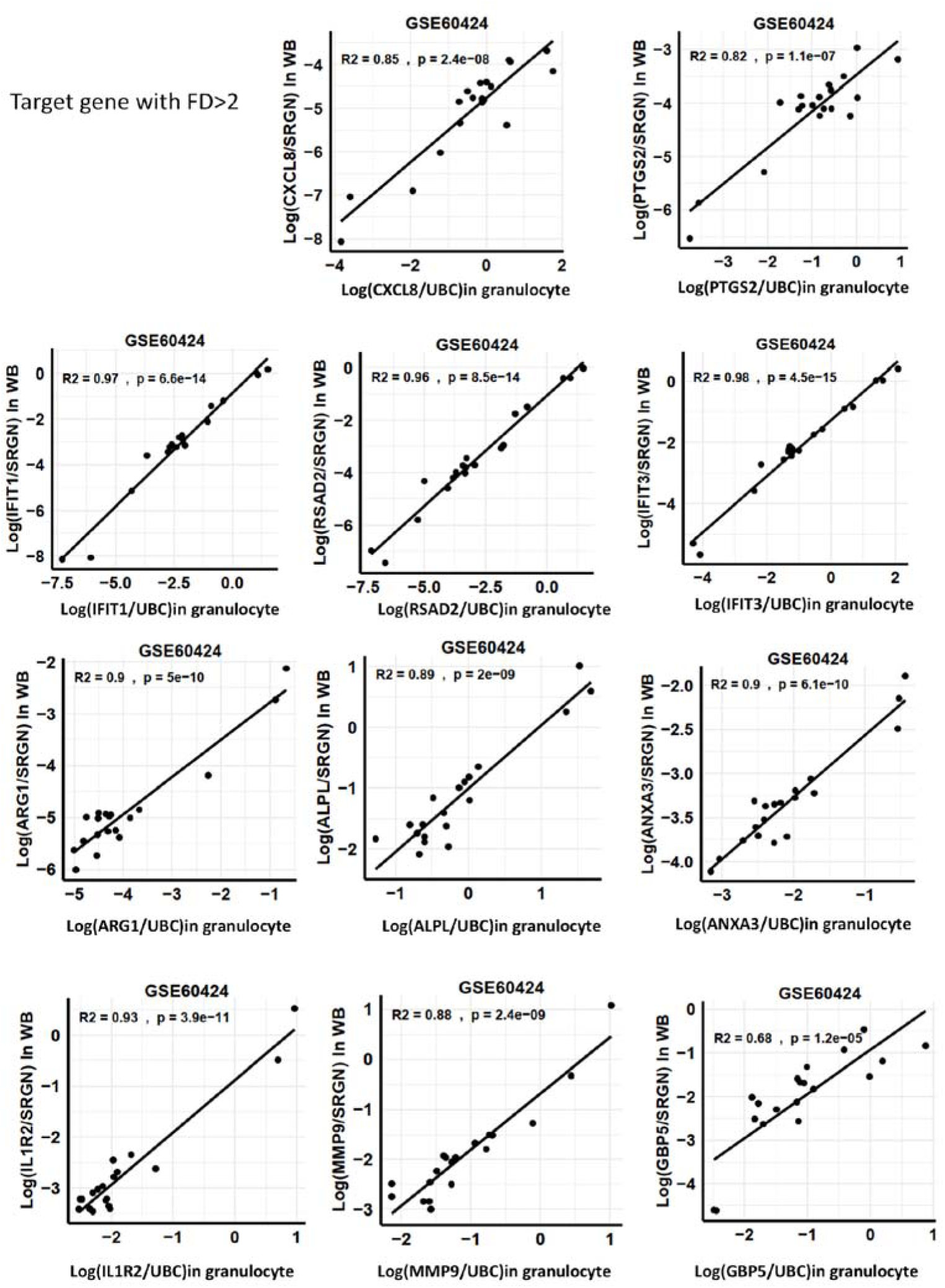
The correlation between DIRECT LS-TA RBB in WB (Y axis, DIRECT LS-TA RBB, target gene as numerator and *SRGN* as denominator) vs gene expression in manually purified granulocytes as the Gold Standard (X axis, target gene expression in purified granulocytes normalised to *UBC*).

Table 1 lists 64 validated granulocyte-informative genes showing excellent correlation between DIRECT LS-TA RBB and gold standard expression level in purified granulocytes. *SRGN* was used as the denominator (granulocyte-informative reference gene) in this DIRECT LS-TA RBB analysis. Bulk WB expression data from GSE60424 was used. All Pearson’s Correlation coefficients (R) are above 0.8 and statistically significant (P<0.001).

### Granulocyte DIRECT LS-TA RBBs useful for the detection of bacterial infection

One RNA sequencing dataset was used as a discovery dataset to identify DIRECT LS-TA RBB that can serve as biomarkers of host response to bacterial infection. GSE261482, an RNA sequencing dataset with 40 bacterial patients and 38 healthy controls, was used as a discovery dataset. Figure 4 shows the volcano plot of the between-group difference in DIRECT LS-TA RBB listed in Table 1. There were 12 up-regulated DIRECT LS-TA RBBs, including *ANXA3/SRGN* and *ARG1/SRGN*. They were chosen for subsequent meta-analysis. On the other hand, RBBs of other genes (27 genes) were suppressed, like *CCN3/SRGN* and *CCR3/SRGN*. However, as we are more interested in stimulated gene biomarkers, they are not further evaluated here. Of note are two interferon-stimulated genes and their RBBs, *IFIT1/SRGN* and *RSAD2/SRGN*, which showed either no change or slight suppression during bacterial infection. Other upregulated genes included *S100A9* but they are considered cell-type markers, so they are not further analysed here.

**Figure 4.**
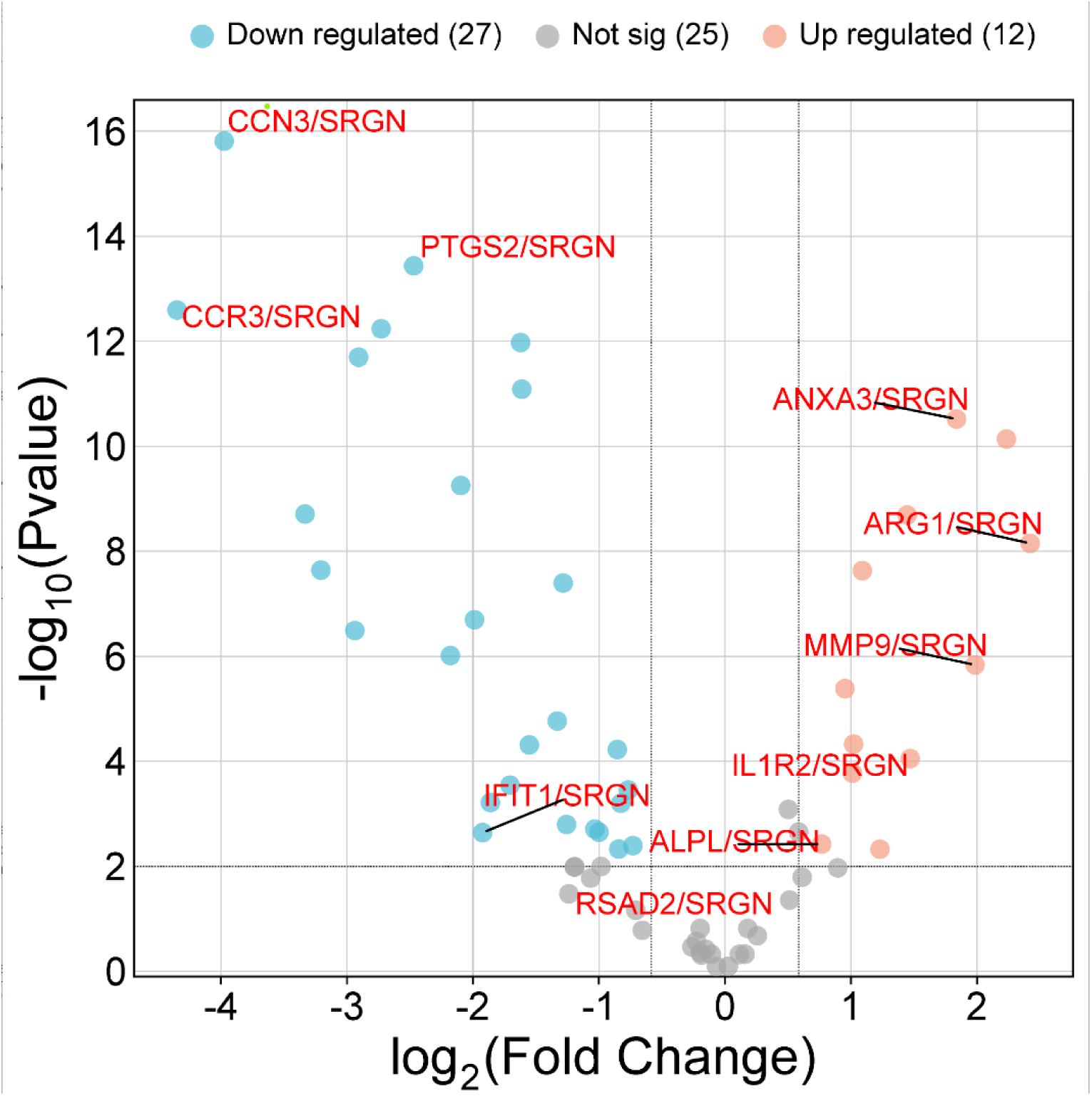
Volcano plot of changes in DIRECT LS-TA RBB in GSE261482 listed in Table 1.

The 5 up-regulated DIRECT LS-TA measured in whole blood samples from patients (labelled as DB) and controls (labelled as C) are shown as a Box plot in Figure 5A. *ANXA3/SRGN* had the best AUC (0.93) in receiver operating characteristic (ROC) analysis (Figure 5B).

**Figure 5A.**
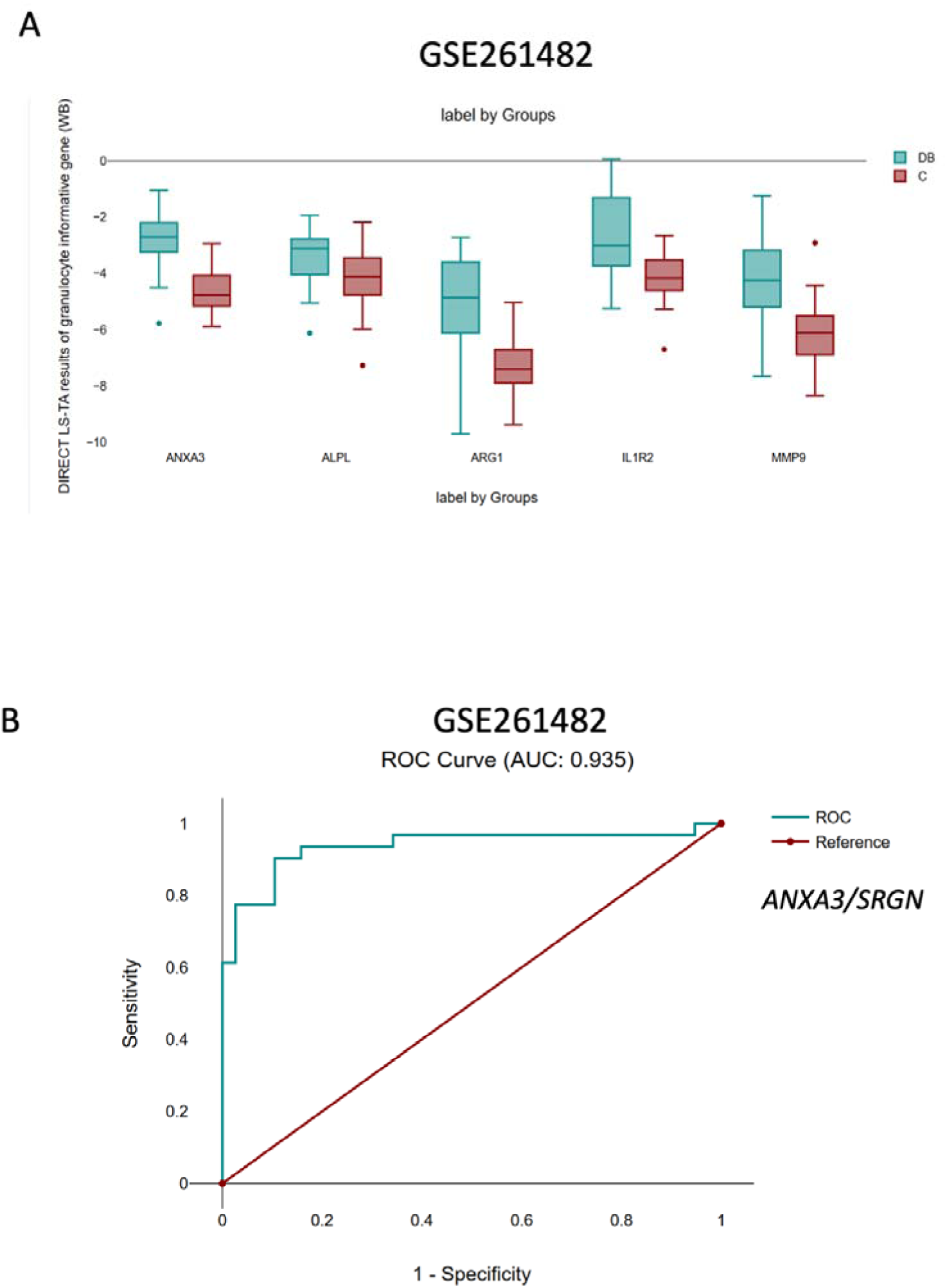
Box plot of 5 up-regulated DIRECT LS-TA from patients (DB) and healthy controls (C) in GSE261482. Figure 5B. ROC curve of *ANXA3/SRGN* in GSE261482.

### Meta-analysis for bacterial infection RBBs

These five RBB *ALPL/SRGN, ANXA3/SRGN, ARG1/SRGN, IL1R2/SRGN and MMP9/SRGN* were further analysed by meta-analysis (Figure 6).

**Figure 6.**
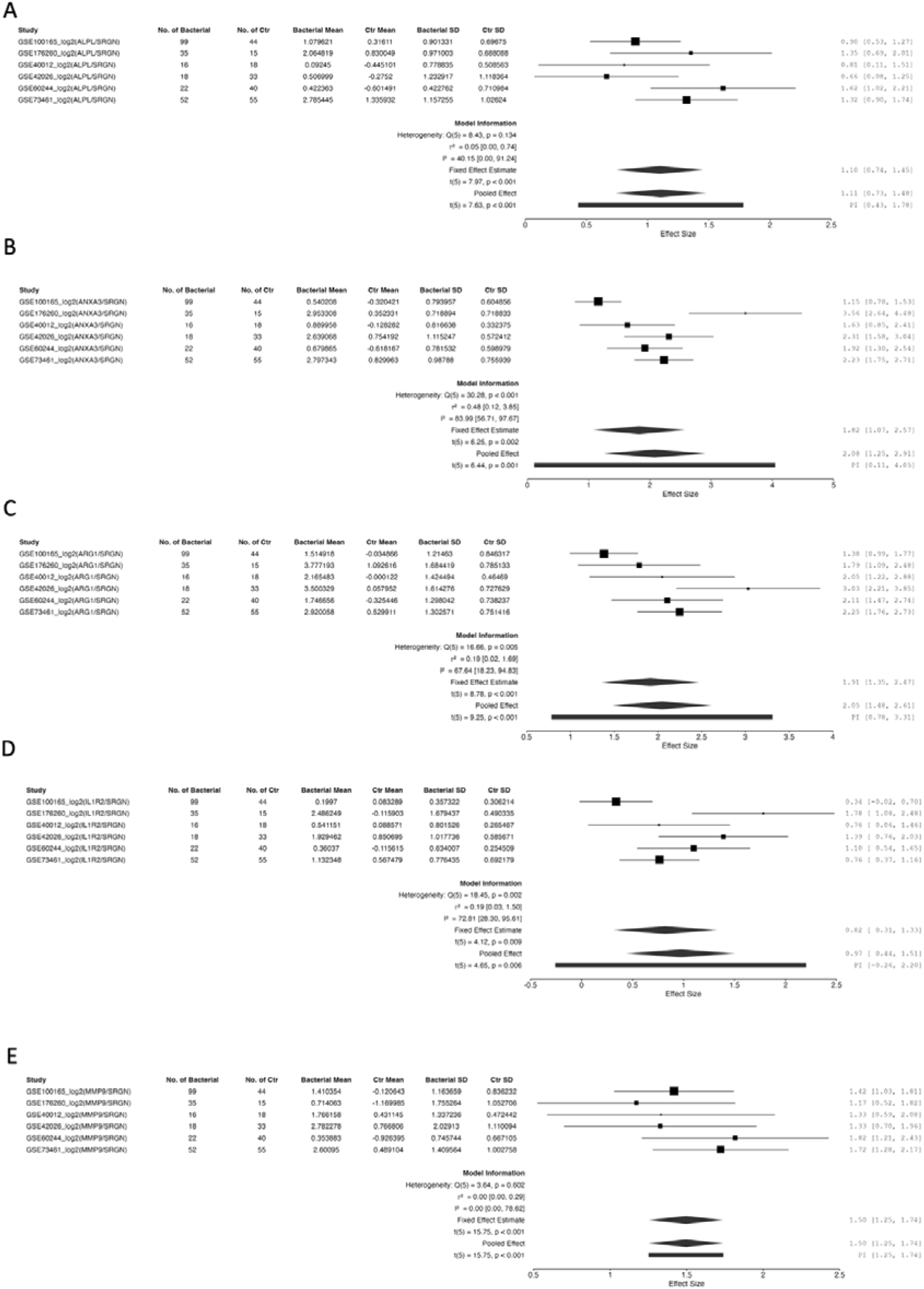
Meta-analysis of (A) *ALPL/SRGN*, (B) *ANXA3/SRGN,* (C)*ARG1/SRGN,* (D)*IL1R2/SRGN and* (E)*MMP9/SRGN* in the detection of bacterial infection.

The ROC curves for these 5 DIRECT LS-TA RBBs in detecting bacterial infection across various datasets are shown in Supplementary Figure 1.

### Granulocyte DIRECT LS-TA RBBs useful for the detection of viral infection

Again, an RNA-sequencing dataset (GSE157240) was used as a discovery dataset to identify useful granulocyte DIRECT LS-TA RBB in detecting the host response to viral infection. GSE157240 included data of patients infected by different types of viruses. Figure 7A shows the volcano plot of DIRECT LS-TA RBBs in the detection of viral infection (all types of viral infection as one group). Three RBBs stand out, *RSAD2/SRGN*, *IFIT1/SRGN* and *IFIT3/SRGN*. *IFIT1* and *IFIT3* are highly correlated in terms of expression; therefore, their performance in DIRECT LS-TA RBB are also very similar. Figure 7B shows the classification performance of 3 example DIRECT LS-TA RBBs in the detection of viral infection.

**Figure 7.**
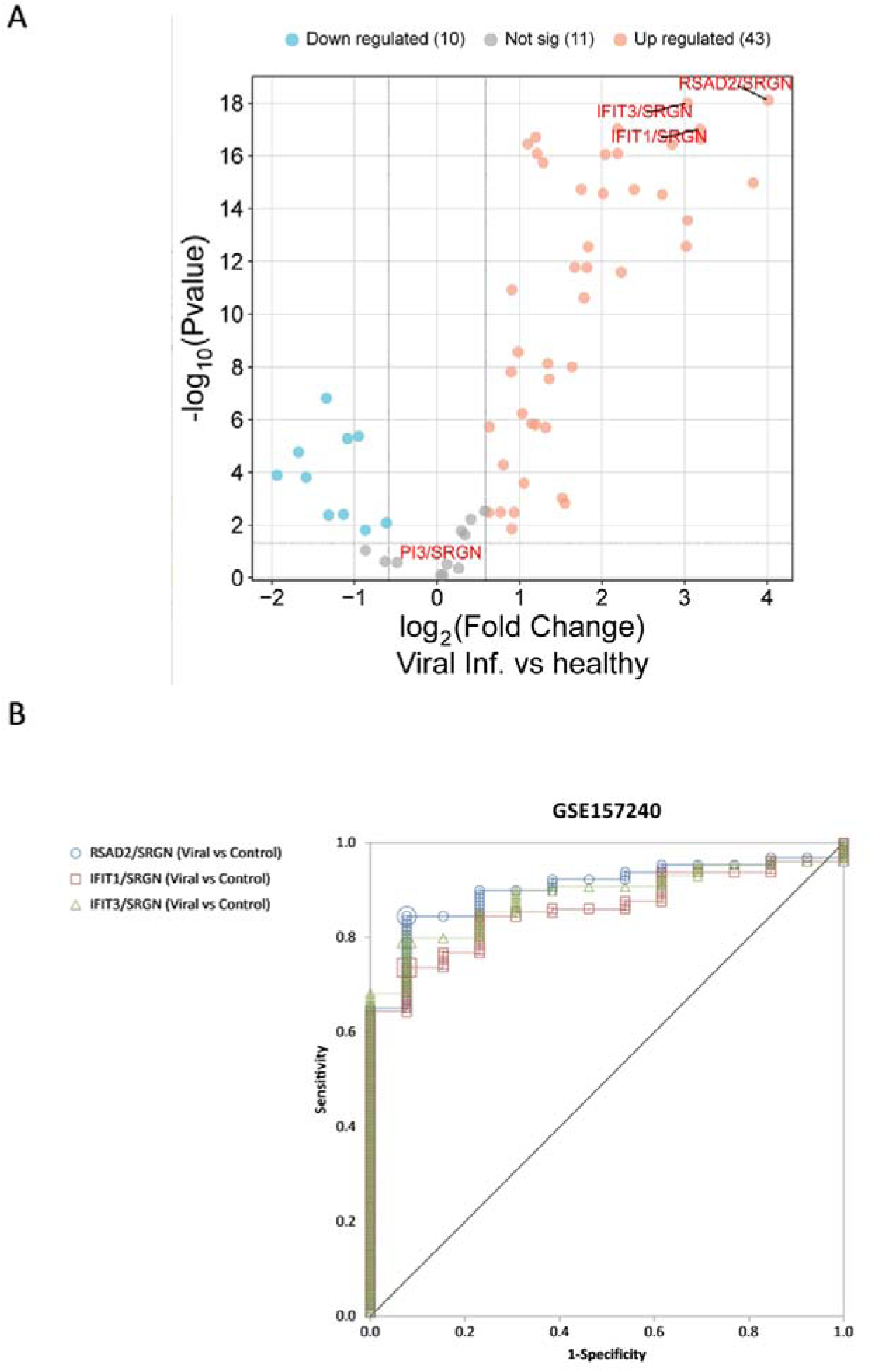
(A) Volcano plot of DIRECT LS-TA RBBs in the detection of viral infection in GSE157240. All types of viral infection are grouped as one group. (B) Classification performance of *RSAD2/SRGN*, *IFIT1/SRGN* and *IFIT3/SRGN* in detection of viral infection in GSE157240.

The 5 up-regulated DIRECT LS-TA (*RSAD2/SRGN, IFIT1/SRGN, IFIT3/SRGN, HERC5/SRGN and SERPING1/SRGN)* measured in whole-blood samples from patients infected with various viruses and controls are shown as a Box plot in Figure 8. *RSAD2/SRGN* had the best AUC (0.9) in ROC analysis (Figure 7B). Three RBBs (*RSAD2/SRGN*, *IFIT1/SRGN* and *IFIT3/SRGN*) are highly correlated, for example, the correlation between *IFIT1/SRGN* and *RSAD2/SRGN* is shown in Figure 8B.

**Figure 8.**
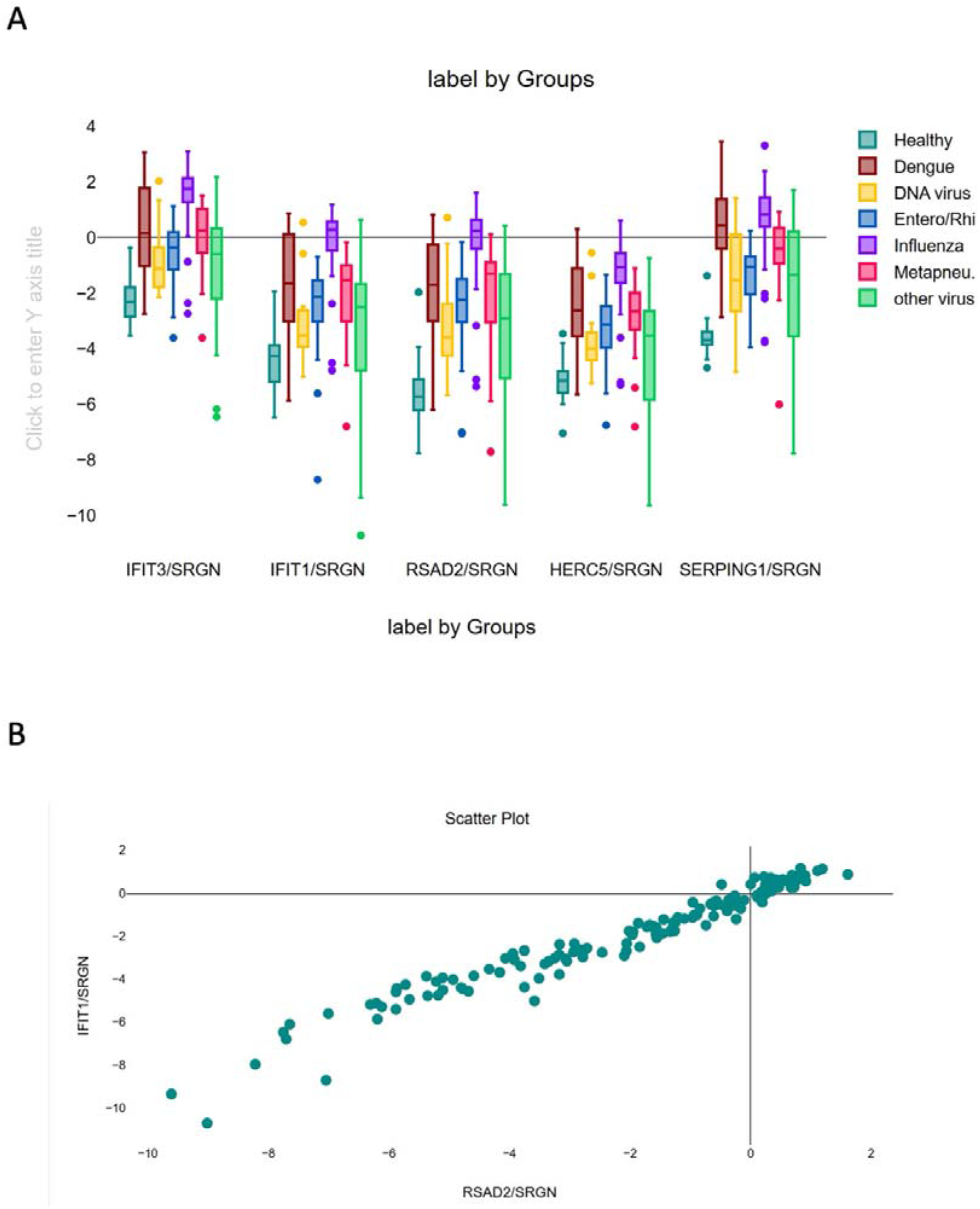
(A) Box plot of RSAD2/SRGN, IFIT1/SRGN and IFIT3/SRGN in different groups of patients in GSE157240. (B) Correlation of IFIT1/SRGN and RSAD2/SRGN in patients in GSE157240.

### Meta-analysis for viral infection RBBs

These three RBB *RSAD2/SRGN, IFIT1/SRGN, IFIT3/SRGN* were further analysed by meta-analysis (Figure 9).

**Figure 9.**
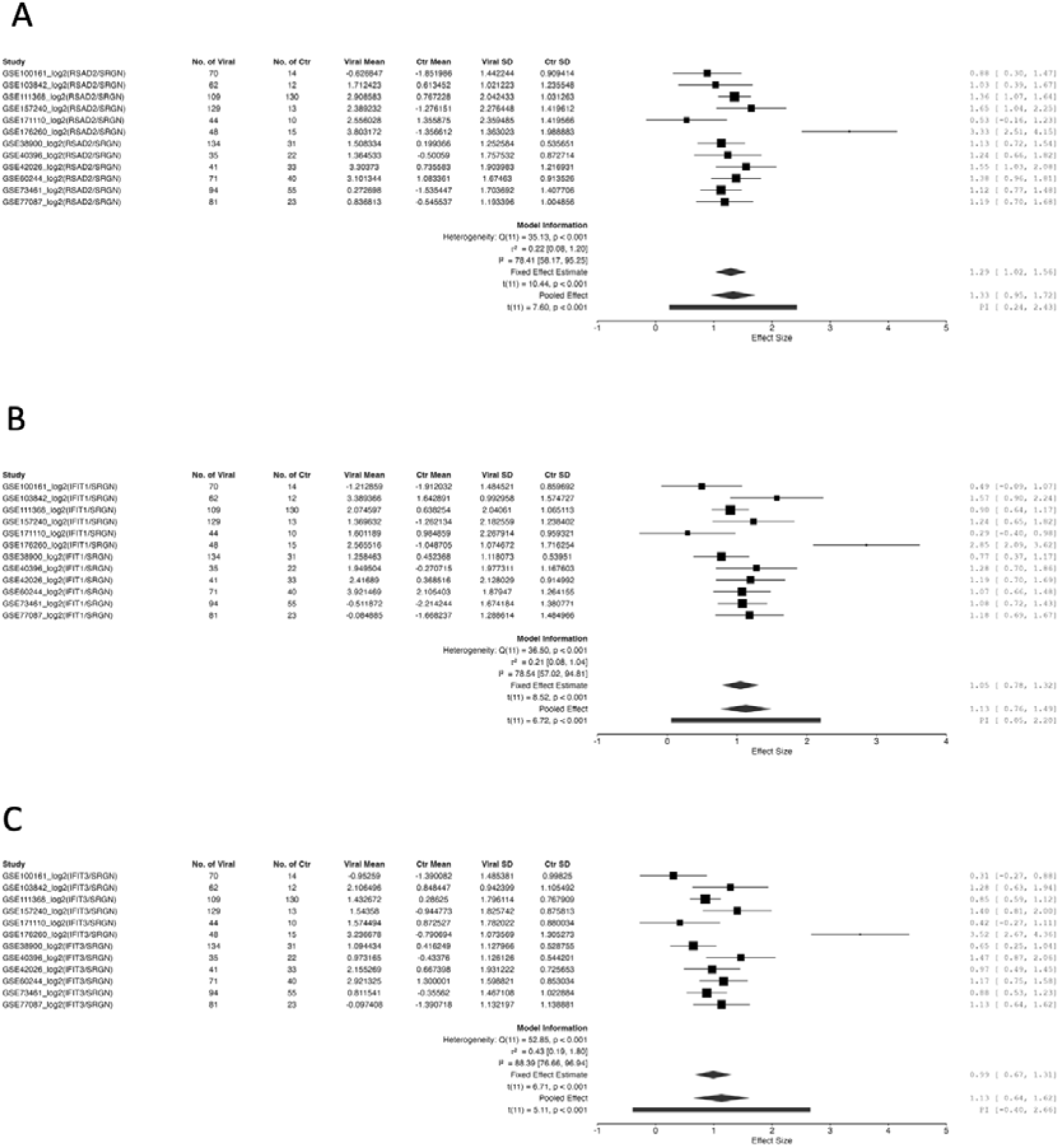
Meta-analysis of (A) *RSAD2/SRGN*, (B) *IFIT1/SRGN and* (C) *IFIT3/SRGN* in the detection of viral infection.

### Evaluation of real-world Classification Performance of RBB: double-ROC in datasets with both bacterial and viral infection patients

In a clinic setting, a febrile patient comes to medical attention or the Emergency Room (ER). So the real-world clinical question is whether the patient gets a bacterial or viral infection. Most public datasets only compare one type of infection against healthy control individuals. Information from such datasets does not directly answer the clinical question. Only datasets that include data from both patient groups (even better, including the control group) are most valuable for evaluation.

Three RNA-sequencing datasets with data from both bacterial and viral infection patients are present: GSE161731, GSE211567, GSE176260, and the following three microarray datasets: GSE73461, GSE42026, GSE60244. ROC was performed to compare between viral and bacterial patients. Previously reported DIRECT LS-TA RBB of monocyte-informative genes (*VNN1/PSAP* as bacterial biomarker and *IFI27/PSAP* as viral biomarker) were also included in the same ROC graph to show the relative performance of each RBB (Figure 10 and Suppl Figure 2).

**Figure 10.**
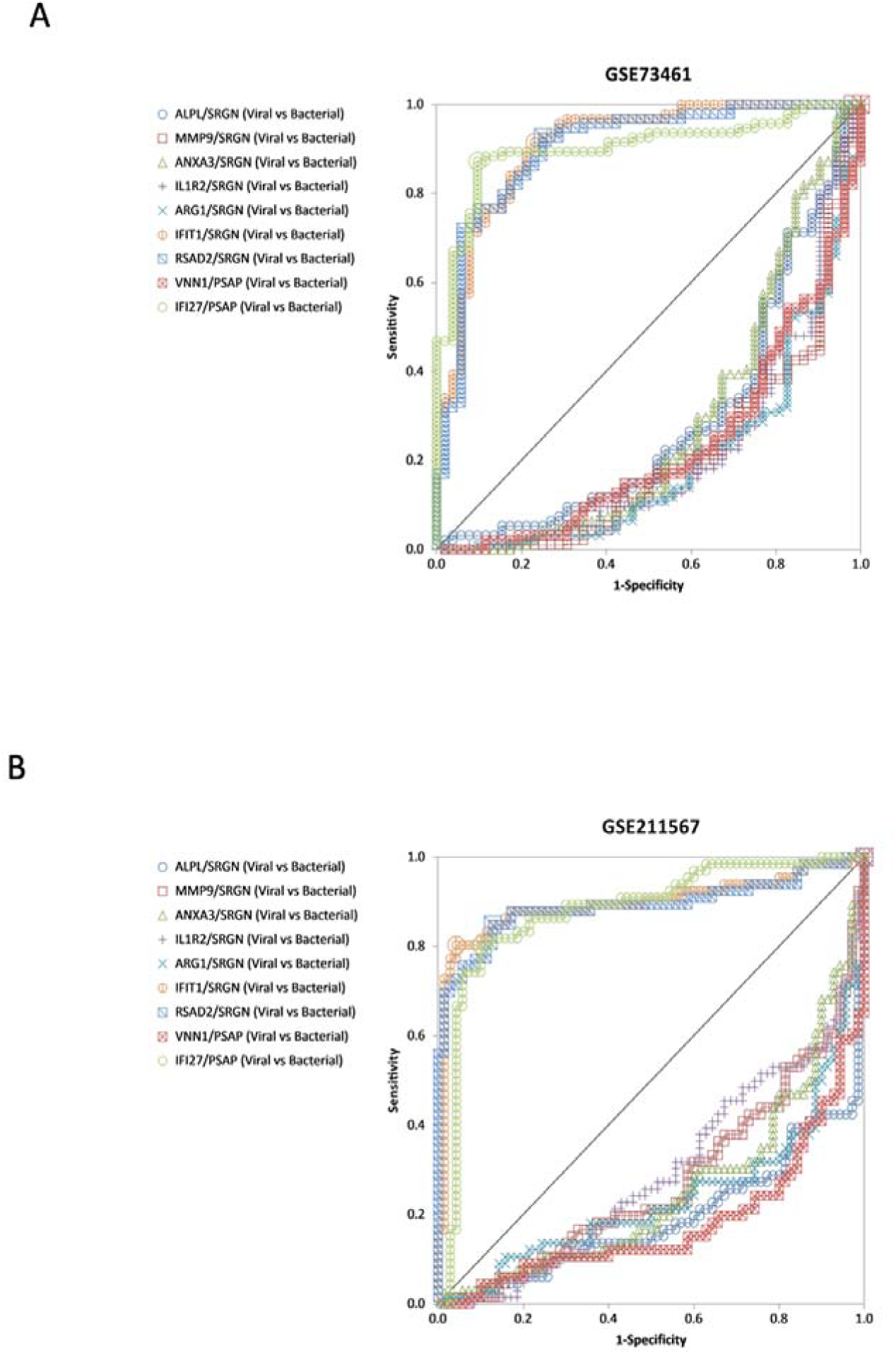
Receiver operating characteristic (ROC) curve of biomarkers in GSE73461 (A) and (B) GSE211567. Double ROC curve for two sets of RBB is shown. RBB for the detection of viral infection are shown on the left side, and those for the detection of bacterial infection are shown on the right side. The best-performing ROC (AUC=1) for viral infection detection will be a ROC curve close to the upper-left corner, and that for RBBs detecting bacterial infection will be close to the lower-right corner.

For example, GSE73461 has data from patients with different types of infection. WB gene expression results from groups of patients were analysed by the DIRECT LS-TA method (Definitive Bacterial infection: N=52, and Definitive Viral infection: N=94). The results are shown in Figure 10 as a double-ROC curve plot. The ROC on the left side shows the performance of RBBs in detecting viral infection, and the ROC on the right side shows their performance in detecting bacterial infection. It is apparent that ROCs of RBBs to viral detection perform better than those of RBBs to detect bacterial infection. On the left side of Figure 10, the ROC curves for RBBs (e.g. *RSAD2/SRGN*) detecting of viral infection is much closer to the top left corner, whereas RBBs (e.g. *IL1R2/SRGN*) for detection of bacterial infection, shown on the right side of the figure, are further away from the lower right corner (perfect point).

Performance of RBBs differs among datasets. For the detection of viral infection, both granulocyte (e.g. *RSAD2/SGRN*) and monocyte direct LS-TA RBB (e.g. IFI27/PSAP) performed well. For detection of bacterial infection, these two Direct LS-TA RBB appear superior to the rest (*IL1R2/SRGN, VNN1/PSAP*). In general, combining a granulocyte DIRECT LS-TA RBB (e.g. *RSAD2/SGRN*) together with a monocyte DIRECT LS-TA RBB (e.g. *VNN1/PSAP*) or vice versa appears to provide the optimal diagnostic yield. This is the advantage of using a single cell-type biomarker as Direct LS-TA of different cell types can be used together in diagnosis. They also carry a cell-of-origin identity, and a more clearly defined diagnostic algorithm can be entertained.

### Multivariate Classification Approaches with Machine Learning

A multivariate approach using multiple DEGs or RBBs should enhance performance for differentiating diseases. Recently, there are various gene panels proposed to perform this same task. To make a fair comparison, only gene panels with fewer than 6 genes are evaluated [12–16,26]. There is a problem applying the original reported risk model parameters to other datasets as TA results are generated by different analysis platforms. To circumvent the problem, an unsupervised machine learning approach is used to compare gene panels using only the data from the selected genes in each panel. In essence, the information content of the selected genes is compared across different panels. WEKA software provides various unsupervised classification algorithms that extract information from variables using a machine learning approach [27]. Three such methods are used for classification that cover possible underlying latent risk models, both linear and nonlinear ones (Naive Bayes, Support Vector Machine (SVM), and random forest). Expression data up to 6 genes, as prescribed by the gene panels, were entered into WEKA. For the DIRECT LS-TA RBBs, the 2-ratio results (RBBs) are provided directly to WEKA; that is, only 2 variables (RBBs) are derived from 4 genes (2 numerator target genes and 2 denominator genes). (AUC in Table 2 and Specificity and Sensitivity in Supplementary Table 1 and 2 respectively)

**Table 2.**
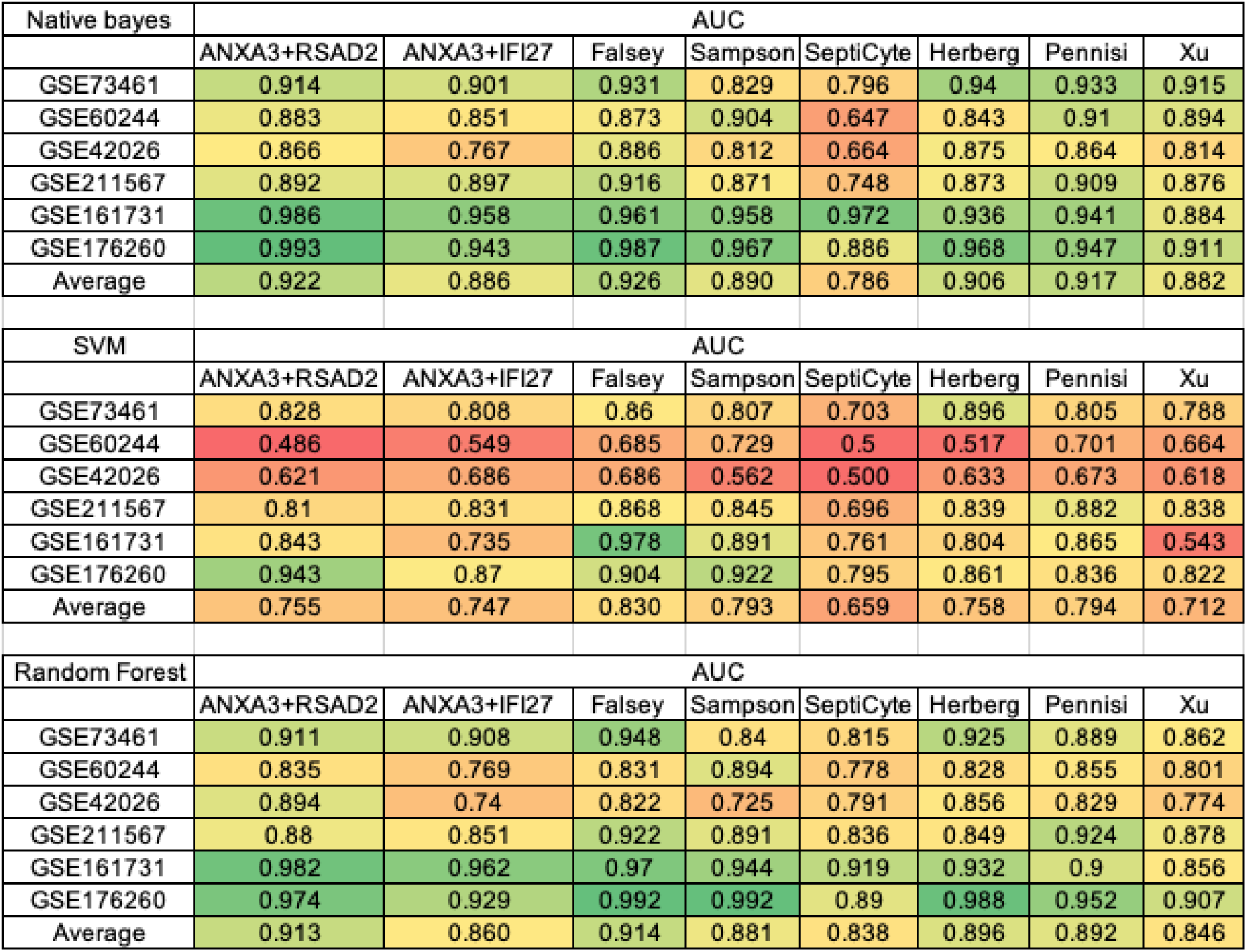
Evaluation of machine learning algorithms on the performance (AUC) of RBB and other gene panels in datasets with both viral and bacterial infection data. *ANXA3*+*RSAD2* : *ANAX3/SRGN*+*RSAD2/SRGN; ANXA3+IFI27: ANXA3/SRGN+IFI27/PSAP;* Falsey: Falsey et al 2025[12]; Sampson: Sampson et al 2017[13]; SeptiCyte: SeptiCyte LAB [26]; Herberg: Herberg et al 2016[14]; Pennisi: Pennisi et al 2021[15]; Xu: Xu et al 2021[16]. List of genes used in each gene panel are listed in Suppl table 3.

## Discussion

Our previous works [20,21] developed the algorithm to obtain monocyte single-cell-type gene expression as a host response to various types of infection. Granulocytes are present in a much higher proportion in the peripheral whole-blood sample; therefore, many more genes (∼10,000 genes) meet the requirement for granulocyte-informative genes. It is widely acknowledged that granulocytes heavily influence bulk gene expression in WB, so immunology researchers often use PBMCs in their studies to avoid the substantial contribution of granulocytes to transcript abundance. There has been no systematic algorithm for evaluating the contribution of granulocytes to the bulk transcript levels in WB. Our in-silico cell-sorting approach addressed this gap and leveraged the high proportion of granulocytes in WB to derive a single-cell-type transcript biomarker for granulocytes [28]. It allows a complete omission of the tedious cell-sorting laboratory procedure to obtain single-cell-type transcriptome biomarkers, which can be used in the triage of febrile illness. It is always preferable to have an assessment of gene expression for a specified single cell type rather than a bulk expression level from a mixture of many cell populations. Direct LS-TA RBB greatly reduces the confounding effect of variation in cell-count proportions across cell types and yields information that can only be obtained through tedious and expensive laboratory procedures, such as single-cell RNA sequencing. This in-silico sorting approach firstly identifies a list of granulocyte-informative genes which can be used in the subsequent development of biomarkers. These genes are produced predominantly (≥50%) by only a single cell type (granulocytes in this example) in the whole blood.

Typically, granulocytes account for 50-70% of leukocytes in WB, with more than 90% being neutrophils; therefore, granulocytes and neutrophils are used interchangeably in this study. Based on the formula and example shown in Figure 1, genes even expressed at a slightly lower level in purified granulocytes than other cell-types in the WB could be qualified as a granulocyte-informative gene. We assumed 50% granulocyte cell count in our algorithm. At this 50% granulocyte count level, a fold difference (FD) value of 1 fold is required for granulocyte information genes. That is, any gene expressed at the same TA per cell (in laboratory practice, it can also be normalised against a housekeeping gene, like UBC) in manually purified granulocytes and in the WB sample fulfils the definition. Under this condition, for at least 10,000 genes, their mRNA transcripts present in WB are produced predominantly by granulocytes alone. Only genes that had excellent correlation with DIRECT LS-TA are listed in Table 1, and they have a virtually perfect correlation between DIRECT LS-TA RBB and TA of purified granulocytes (R>0.9).

Then, these DIRECT LS-TA RBBs were evaluated in public datasets of patients with viral or bacterial infections for their triage performance. The RBB is a ratio of a granulocyte-informative target gene to a granulocyte-informative reference gene (e.g. SRGN). Therefore, for the first time, a gene expression RBB assay can be used to represent single-cell-type gene expression in granulocytes and to reveal stimulation or suppression of the target gene due to pathology, such as the host response to infection. In the case of RBBs in viral infection, the granulocyte target genes are interferon-stimulated genes (ISGs) such as *IFIT1* and *RSAD2.* By using Direct LS-TA RBB of ISG, we avoid the confounding effect of variation in cell-type proportions when interpreting granulocyte gene expression results. Another advantage of the DIRECT LS-TA approach is that the in-silico cell-sorting is done early during the assay development stage, and the shortlisted DIRECT LS-TA can be used subsequently in clinical practice. The RBBs can be easily measured by either digital or quantitative PCR using WB (whole blood) samples. Samples collected in RNA fixative, such as PAX-tube, are compatible with our approach.

Take *RSAD2,* for example, the granulocyte single-cell-type gene expression of *RSAD2* is represented by a ratio of the transcript abundance of *RSAD2* to that of *SRGN*. *SRGN* is the granulocyte single-cell-type reference gene as stipulated by the in silico cell-sorting approach. As revealed in Figure 2, DIRECT LS-TA results of *RSAD2* (results obtained from WB specimens shown in the Y axis) are highly correlated with *RSAD2* expression in purified granulocytes obtained from the same individual (R2∼0.96, P=8.5e-14). Similar correlations are observed for other ISGs (e.g., *IFIT1* and *IFIT3; results not shown*). Therefore, granulocyte single-cell-type gene expression of these ISGs can be reliably estimated directly from a WB specimen without the need to manually separate granulocytes. ISGs play important roles during host response during viral infection, and they also serve as good biomarkers of viral infection[29–31].

For host response during bacterial infection, DIRECT LS-TA RBB of 5 genes are shortlisted for evaluation in Table 1. The most definitive test is one that mirrors a real-world scenario in which a febrile patient presents to the ER for consultation. The first clinical question is whether it is caused by a viral or bacterial pathogen. Host response biomarkers have been sought to answer this question. Therefore, 6 selected datasets containing patients with viral and bacterial infections were further analysed. Both ROC curves left-upper corner (the typical ROC), which reveals the performance of viral infection detection, and the right lower corner ROC, reveals the performance of bacterial infection detection are important.

Another distinct advantage of using the DIRECT LS-TA assay is that different DIRECT LS-TA assays concerning different cell populations can be used together, such as a granulocyte LS-TA result (e.g. the RBB of *ANXA3/SRGN*) together with a monocyte LS-TA result (e.g. the RBB of *IFI27/PSAP*). Only the measurement of TA of 4 genes is required to generate these 2 ratios reflecting target gene expression in 2 different cell types. The outstanding performance of such a combination was demonstrated in an unsupervised classification test against other gene panels derived from traditional DEG algorithms which did not take into account of cell-type proportions of various cell-types in WB.

## Acknowledgement

During the preparation of this work, the authors used Grammarly for English editing and correction of grammatical usage. The work described in this paper was supported by a grant from Research Grants Council of Hong Kong Special Administrative Region, China (Project No. 14104625) and the Co-funding Mechanism on Joint Laboratories with the CAS (JLFS/M-403/24).

