## Supplementary material for "In-silico cell sorting revealed granulocyte-specific single-cell-type gene expression from peripheral blood bulk expression data and its application as host response biomarkers to discriminate bacterial and viral infections"

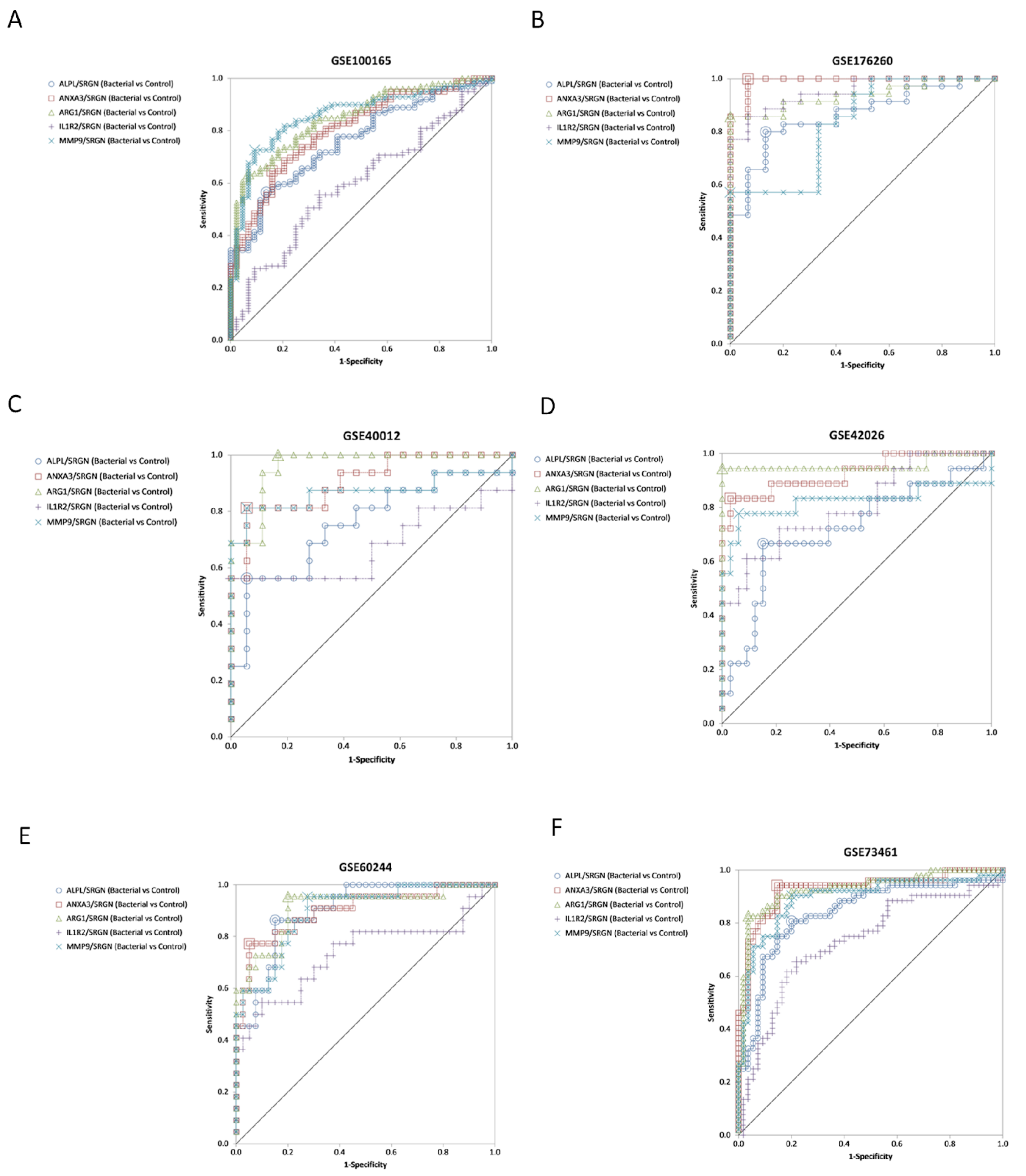


**Supplementary Figure 1. Receiver operating characteristic (ROC) curve of DIRECT LS-TA RBBs *ALPL/SRGN, ANXA3/SRGN, ARG1/SRGN, IL1R2/SRGN and MMP9/SRGN* in the detection of bacterial infection in datasets (A)GSE100165, (B)GSE176260, (C)GSE40012, (D)GSE42026, (E)GSE60244 and (F)GSE73461.**

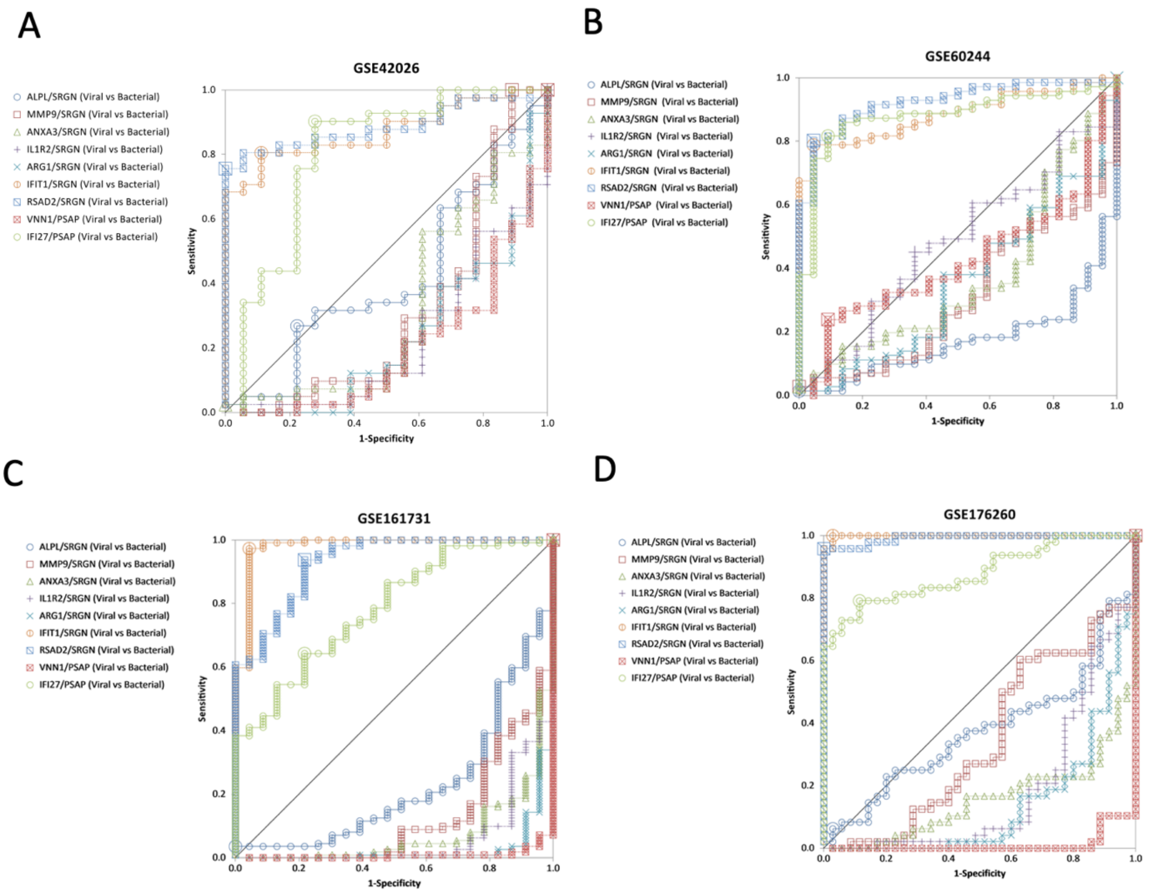


**Supplementary Figure 2. Receiver operating characteristic (ROC) curve of biomarkers in (A) GSE42026, (B) GSE60244, (C) GSE161731 and (D) GSE176260**

| Native bayes | Specificity | | | | | | | |
|---|---|---|---|---|---|---|---|---|
| | ANXA3+RSAD2 | ANXA3+IFI27 | Falsey | Sampson | SeptiCyte | Herberg | Pennisi | Xu |
| GSE73461 | 0.858 | 0.863 | 0.863 | 0.792 | 0.714 | 0.871 | 0.817 | 0.818 |
| GSE60244 | 0.793 | 0.696 | 0.626 | 0.821 | 0.22 | 0.777 | 0.831 | 0.79 |
| GSE42026 | 0.685 | 0.615 | 0.777 | 0.632 | 0.615 | 0.716 | 0.731 | 0.639 |
| GSE211567 | 0.81 | 0.825 | 0.86 | 0.838 | 0.696 | 0.841 | 0.848 | 0.83 |
| GSE161731 | 0.853 | 0.743 | 0.925 | 0.815 | 0.746 | 0.818 | 0.818 | 0.495 |
| GSE176260 | 0.933 | 0.865 | 0.9 | 0.908 | 0.832 | 0.867 | 0.899 | 0.867 |
| Average | 0.822 | 0.768 | 0.825 | 0.801 | 0.637 | 0.815 | 0.824 | 0.740 |

| SVM | Specificity | | | | | | | |
|---|---|---|---|---|---|---|---|---|
| | ANXA3+RSAD2 | ANXA3+IFI27 | Falsey | Sampson | SeptiCyte | Herberg | Pennisi | Xu |
| GSE73461 | 0.8 | 0.78 | 0.851 | 0.785 | 0.639 | 0.882 | 0.767 | 0.76 |
| GSE60244 | 0.23 | 0.355 | 0.564 | 0.608 | 0.237 | 0.293 | 0.595 | 0.554 |
| GSE42026 | 0.531 | 0.592 | 0.592 | 0.43 | 0.305 | 0.538 | 0.584 | 0.507 |
| GSE211567 | 0.858 | 0.831 | 0.868 | 0.845 | 0.693 | 0.84 | 0.882 | 0.838 |
| GSE161731 | 0.746 | 0.566 | 0.964 | 0.82 | 0.603 | 0.675 | 0.782 | 0.243 |
| GSE176260 | 0.934 | 0.872 | 0.892 | 0.916 | 0.782 | 0.843 | 0.817 | 0.901 |
| Average | 0.683 | 0.666 | 0.789 | 0.734 | 0.543 | 0.679 | 0.738 | 0.634 |

| Random Forest | Specificity | | | | | | | |
|---|---|---|---|---|---|---|---|---|
| | ANXA3+RSAD2 | ANXA3+IFI27 | Falsey | Sampson | SeptiCyte | Herberg | Pennisi | Xu |
| GSE73461 | 0.821 | 0.838 | 0.857 | 0.736 | 0.714 | 0.883 | 0.777 | 0.739 |
| GSE60244 | 0.651 | 0.603 | 0.585 | 0.689 | 0.453 | 0.484 | 0.63 | 0.509 |
| GSE42026 | 0.777 | 0.6 | 0.615 | 0.47 | 0.577 | 0.716 | 0.547 | 0.462 |
| GSE211567 | 0.821 | 0.812 | 0.88 | 0.822 | 0.769 | 0.815 | 0.866 | 0.836 |
| GSE161731 | 0.848 | 0.84 | 0.89 | 0.743 | 0.704 | 0.815 | 0.817 | 0.59 |
| GSE176260 | 0.982 | 0.849 | 0.957 | 0.941 | 0.822 | 0.9 | 0.848 | 0.848 |
| Average | 0.817 | 0.757 | 0.797 | 0.734 | 0.673 | 0.769 | 0.748 | 0.664 |

**Supplementary Table 1. Evaluation of machine learning algorithms on the performance (Specificity) of RBB and other gene panels in datasets with both viral and bacterial infection data.** *ANXA3+RSAD2* : *ANAX3/SRGN+RSAD2/SRGN; ANXA3+IFI27: ANXA3/SRGN+IFI27/PSAP;* Falsey: Falsey et al 2025[1]; Sampson: Sampson et al 2017[2]; SeptiCyte: SeptiCyte LAB [3]; Herberg: Herberg et al 2016[4]; Pennisi: Pennisi et al 2021[5]; Xu: Xu et al 2021[6]. List of genes used in each gene panel are listed in Suppl table 3.

| Native bayes | Sensitivity | | | | | | | |
|---|---|---|---|---|---|---|---|---|
| | ANXA3+RSAD2 | ANXA3+IFI27 | Falsey | Sampson | SeptiCyte | Herberg | Pennisi | Xu |
| GSE73461 | 0.884 | 0.877 | 0.877 | 0.795 | 0.795 | 0.89 | 0.856 | 0.842 |
| GSE60244 | 0.839 | 0.828 | 0.806 | 0.828 | 0.71 | 0.785 | 0.86 | 0.828 |
| GSE42026 | 0.78 | 0.763 | 0.847 | 0.729 | 0.763 | 0.78 | 0.814 | 0.746 |
| GSE211567 | 0.809 | 0.824 | 0.86 | 0.838 | 0.699 | 0.838 | 0.846 | 0.831 |
| GSE161731 | 0.956 | 0.926 | 0.97 | 0.941 | 0.941 | 0.956 | 0.956 | 0.896 |
| GSE176260 | 0.94 | 0.867 | 0.916 | 0.916 | 0.843 | 0.892 | 0.904 | 0.892 |
| Average | 0.868 | 0.848 | 0.879 | 0.841 | 0.792 | 0.857 | 0.873 | 0.839 |

| SVM | Sensitivity | | | | | | | |
|---|---|---|---|---|---|---|---|---|
| | ANXA3+RSAD2 | ANXA3+IFI27 | Falsey | Sampson | SeptiCyte | Herberg | Pennisi | Xu |
| GSE73461 | 0.856 | 0.836 | 0.87 | 0.829 | 0.767 | 0.911 | 0.842 | 0.815 |
| GSE60244 | 0.742 | 0.742 | 0.806 | 0.849 | 0.763 | 0.742 | 0.806 | 0.774 |
| GSE42026 | 0.712 | 0.78 | 0.78 | 0.695 | 0.695 | 0.729 | 0.763 | 0.729 |
| GSE211567 | 0.86 | 0.831 | 0.868 | 0.846 | 0.699 | 0.838 | 0.882 | 0.838 |
| GSE161731 | 0.941 | 0.904 | 0.993 | 0.963 | 0.919 | 0.933 | 0.948 | 0.844 |
| GSE176260 | 0.952 | 0.867 | 0.916 | 0.928 | 0.807 | 0.88 | 0.855 | 0.843 |
| Average | 0.844 | 0.827 | 0.872 | 0.852 | 0.775 | 0.839 | 0.849 | 0.807 |

| Random Forest | Sensitivity | | | | | | | |
|---|---|---|---|---|---|---|---|---|
| | ANXA3+RSAD2 | ANXA3+IFI27 | Falsey | Sampson | SeptiCyte | Herberg | Pennisi | Xu |
| GSE73461 | 0.863 | 0.863 | 0.897 | 0.788 | 0.795 | 0.897 | 0.815 | 0.808 |
| GSE60244 | 0.785 | 0.731 | 0.774 | 0.806 | 0.753 | 0.753 | 0.817 | 0.731 |
| GSE42026 | 0.847 | 0.729 | 0.763 | 0.644 | 0.746 | 0.78 | 0.678 | 0.627 |
| GSE211567 | 0.824 | 0.816 | 0.882 | 0.824 | 0.772 | 0.816 | 0.868 | 0.838 |
| GSE161731 | 0.933 | 0.911 | 0.97 | 0.926 | 0.904 | 0.941 | 0.948 | 0.852 |
| GSE176260 | 0.976 | 0.867 | 0.952 | 0.94 | 0.819 | 0.916 | 0.85 | 0.855 |
| Average | 0.871 | 0.820 | 0.873 | 0.821 | 0.798 | 0.851 | 0.829 | 0.785 |

**Supplementary Table 2. Evaluation of machine learning algorithms on the performance (Sensitivity) of RBB and other gene panels in datasets with both viral and bacterial infection data.** *ANXA3+RSAD2* : *ANAX3/SRGN+RSAD2/SRGN; ANXA3+IFI27: ANXA3/SRGN+IFI27/PSAP;* Falsey: Falsey et al 2025[1]; Sampson: Sampson et al 2017[2]; SeptiCyte: SeptiCyte LAB [3]; Herberg: Herberg et al 2016[4]; Pennisi: Pennisi et al 2021[5]; Xu: Xu et al 2021[6]. List of genes used in each gene panel are listed in Suppl table 3.

| Model | Genes | Reference |
|---|---|---|
| ANXA3+RSAD2 | *ANXA3/SRGN, RSAD2/SRGN* | NA |
| ANXA3+IFI27 | *ANXA3/SRGN, IFI27/PSAP* | NA |
| Falsey | *ITGA7, IFI27, FAM20A, ITGB4* | [1] |
| Sampson | *ISG15, IL16, OASL, ADGRE5* | [2] |
| SeptiCyte | *PLAC8, PLA2G7, LAMP1, CEACAM4* | [3] |
| Herberg | *FAM89A, IFI44L* | [4] |
| Pennisi | *ADGRE1, IFI44L* | [5] |
| Xu | *PI3, IFI44L* | [6] |

**Supplementary Table 3. List of genes in gene panel used in WEKA analysis.**
